# H3K27M Mutation Reshapes the Antigenic Landscape in Diffuse Midline Glioma

**DOI:** 10.1101/2025.04.07.647502

**Authors:** Tima Shamekhi, Bijun Zeng, Claire Xin Sun, Paul Daniel, Terry C.C Lim Kam Sian, Gabriel Goncalves, Grace Huang, Farnaz Fahimi, Nivedhitha Selvakumar, Erwin Tanuwidjaya, Isaac Woodhouse, Bettina Kritzer, Ralf B. Schittenhelm, Roberta Mazzieri, Jason E. Cain, Javad Nazarin, Jordan R. Hansford, Ron Firestein, Riccardo Dolcetti, Pouya Faridi

## Abstract

**Simple Summary:** Diffuse midline gliomas (DMGs) are aggressive childhood brain tumours with no effective treatments. Over 80% of cases carry the histone H3K27M mutation, which alters chromatin structure and gene regulation and induce tumours growth. Immunotherapy, which uses the body’s immune system to fight cancer, relies on tumour-specific antigen molecules that immune cells can recognise. However, because DMGs have few mutations, finding suitable antigens for therapeutic purposes has been challenging. In this study, we investigated how the H3K27M mutation affects tumour antigen presentation in DMG. Using patient-derived DMG models, we found that H3K27M alters the landscape of antigens displayed on tumour cell surface, creating unique immune targets. We identified six immunogenic peptides, that triggered strong T cell responses. These antigens were absent when H3K27M was removed, confirming their link to the mutation. Our findings provide a blueprint for developing T cell-based immunotherapies for DMG, offering new hope for targeted treatments against this devastating disease.

**Abstract:** *Background:* Diffuse midline gliomas (DMGs) are among the most aggressive paediatric brain tumours, with the pathognomonic H3K27M mutation present in over 80% of cases. This mutation drives epigenetic dysregulation and transcriptional reprogramming, yet its impact on the tumour antigenic landscape remains poorly understood. Given the low mutational burden of DMG, an expanded search beyond neoantigens to include epigenetically dysregulated tumour-associated antigens (TAAs) is critical for advancing antigen-specific immunotherapies.

*Methods:* To assess how H3K27M influences antigenic landscape of DMG, we performed a comprehensive immunopeptidomic analysis using patient-derived DMG cell line models (SU-DIPG13 and BT245) that harbour the H3K27M mutation and their CRISPR-edited H3K27M-knockout (KO) counterparts. High-resolution mass spectrometry and bioinformatics were employed to define H3K27M-driven changes in the immunopeptidome. Functional T cell assays using HLA-matched healthy donor PBMCs were conducted to evaluate the immunogenicity of H3K27M-associated peptides.

*Results:* Our findings reveal that the H3K27M mutation reshapes the tumour antigenic landscape in a model-specific manner. While H3K27M knockout increased HLA-I expression in SU-DIPG13 but not BT245, immunopeptidomic profiling uncovered distinct shifts in the presentation of tumour-associated peptides, independent of direct effects on antigen processing machinery. Among these, we identified six immunogenic peptides, derived from SLITRK2, PRAME, XKR5, and CBX2, that elicited CD8⁺ T cell responses in *in vitro* functional assays. Notably, PRAME, a well-characterised cancer-testis antigen was confirmed as an H3K27M-associated immunogenic target, reinforcing its therapeutic relevance. Peptides identified exclusively in H3K27M+ cells were absent in KO models, demonstrating a direct link between H3K27M-driven transcriptional dysregulation and tumour antigenicity.

*Conclusions:* This study provides the first systematic assessment of how H3K27M reshapes the antigenic landscape in DMG, uncovering novel, immunogenic tumour-associated peptides that could serve as targets for precision immunotherapy. By demonstrating that H3K27M mutation drives context-dependent antigen presentation, our findings establish a foundation for T cell-based therapies targeting H3K27M-associated antigens. These insights pave the way for next-generation personalised immunotherapies for this otherwise treatment-refractory disease.

*Key Points:* H3K27M mutation induces expression of tumour-associated antigens in DMG H3K27M alters the DMG immunopeptidome without uniformly changing HLA-I levels PRAME- and CBX2-derived peptides are immunogenic and targetable by CD8⁺ T cells

*Importance of Study:* Diffuse midline gliomas (DMGs) are universally fatal paediatric brain tumours with limited treatment options and poor immune visibility. While the H3K27M mutation is a defining hallmark, its impact on tumour immunogenicity remains unclear. This study presents the first comprehensive to explore the effect of H3K27M-mution on the DMG immunopeptidome, revealing six immunogenic peptides derived from epigenetically dysregulated tumour-associated antigens, including SLITRK2, PRAME, XKR5, and CBX2. These antigens elicited CD8⁺ T cell responses, establishing a direct link between H3K27M-driven transcriptional dysregulation and tumour antigenicity. By leveraging these altered antigens, we highlight actionable vulnerabilities for T cell-based immunotherapy.

## 1. Introduction

Diffuse midline gliomas (DMGs) represent one of the most aggressive and lethal forms of paediatric brain tumours, with a median survival rate of 9-11 months (1, 2). The heterozygous somatic histone mutation H3K27M is present in over 80% of DMG cases, especially in diffuse intrinsic pontine gliomas (DIPGs) (3, 4). This mutation affects key histone genes: the *HIST1H3B*/C, which encode the canonical histone H3.1, and *H3F3A,* encoding the variant histone H3.3. The H3.3 accounts for roughly 70% of H3K27M mutations and is correlated with a particularly poor prognosis (5, 6).

Mechanistically, substituting lysine 27 with methionine in histone H3 triggers widespread epigenetic reprogramming. This mutation results in a global loss of H3K27 di- and tri- methylation, increased acetylation, and alterations in three-dimensional chromatin architecture, collectively leading to the dysregulation of DNA methylation and gene expression (7, 8). Although only 3–17% of total cellular histone H3 proteins are mutated, the resulting dominant hypomethylation disrupts transcriptional silencing and promotes the upregulation of proto-oncogenes while repressing genes involved in cellular differentiation (9, 10). Beyond driving oncogenesis, these epigenetic alterations induce a stem cell–like, immune-cold tumour state and modulate antigen presentation and T cell infiltration, thereby facilitating immune evasion (9, 11–16).

The advent of immunotherapy, particularly adoptive T cell–based approaches, offers a promising strategy to overcome the inherent resistance of DMG to conventional therapies by enhancing tumour immunogenicity. Effective anti-tumour immunity is largely mediated by CD8⁺ T cells that recognise tumour antigens presented by major histocompatibility complex (MHC) class I molecules (known as human leukocyte antigens, HLA, in humans). Typically, these antigens are peptide fragments derived from tumour-specific mutations or dysregulated proteins (17, 18). Peptide antigen derived from the H3K27M mutation represent promising targets for peptide–HLA (pHLA)–mediated immunotherapy due to their tumour specificity. However, previous efforts to identify HLA-A*02:01–restricted peptide neoantigens derived from this mutant protein have yielded inconclusive results, underscoring the need to expand the search for other actionable cancer antigens (19–21).

The immune-privileged nature of the brain poses additional challenges for effective immunotherapy in DMG (22). Overcoming these barriers requires a comprehensive understanding of the tumour immunopeptidome to drive the development of targeted therapeutic strategies. To bridge this gap, we performed an in-depth immunopeptidomic analysis to systematically define the landscape of tumour-specific antigens in DMG. Given the inherently low mutational burden of DMG and the challenge of identifying H3K27M- derived neoantigens (9, 20, 23), we expanded our search to include non-mutated but aberrantly expressed antigens influenced by the tumour’s epigenetic dysregulation. Using state-of-the-art mass spectrometry and robust bioinformatic pipelines, we sought to determine how the H3K27M mutation reshapes antigen presentation and whether it gives rise to immunogenic pHLA complexes. To achieve this, we leveraged two patient-derived H3K27M- mutant DMG cell line models (SU-DIPG13 and BT245) and their CRISPR-edited H3K27M- knockout (KO) counterparts (7). These models provide a controlled isogenic background, minimising confounding variables and enabling a direct evaluation of the mutation’s impact. Moreover, we functionally assessed the immunogenicity of the identified pHLA complexes to pinpoint novel actionable targets for T cell–mediated immunotherapy.

By integrating these advanced methodologies, our study provides a comprehensive analysis of H3K27M-mutant DMG, offering crucial insights into how this mutation reshapes the tumour’s antigenic landscape. We identify antigenic changes associated with H3K27M- driven epigenetic dysregulation and the generation of targetable antigens. This work not only enhances our understanding of the immune landscape in DMG but also lays the foundation for novel, personalised immunotherapeutic strategies aimed at overcoming the immune-evasive nature of these highly aggressive tumours.

## 2. Materials and Methods

The experimental procedures were designed to investigate the immunogenic profile of H3K27M-mutated cells by analysing the altered immunopeptidome and antigen presentation using isogenic models of paediatric diffuse midline glioma (pDMG).

### 2.1 Cell culture

Two Patient-derived H3K27M-isogenic (H3K27M and H3K27M-KO) cell line pairs, SU- DIPG13 and BT245 were used in this study. The parental H3K27M-mutant cell lines were originally established at Stanford University (SU-DIPG13) and Boston Children’s Hospital (BT245). Subsequently, the H3K27M-KO variants were generated by Dr. Nada Jabado’s team at McGill University (7) . Cells were maintained in tumour stem medium (TSM). Tumour- base medium (TBM) was prepared by mixing Neurobasal-A and Dulbecco’s Modified Eagle Medium/F-12 (DMEM/F-12) (1:1 ratio) (Gibco), supplemented with 10 mM HEPES, 1 mM sodium pyruvate, 1× MEM non-essential amino acids, 1× GlutaMAX, and 1% penicillin-streptomycin (Gibco). TSM was prepared by supplementing TBM with 1× B27 supplement (minus vitamin A), 20 ng/mL EGF, 20 ng/mL FGF-basic, 10 ng/mL PDGF-AA, 10 ng/mL PDGF-BB, and 2 µg/mL heparin (STEMCELL Technologies). Cells were detached using Accutase (Sigma), washed with 1x PBS, and subsequently pelleted by centrifugation at 2500 × g for 15 min at 4°C. Cell pellets were snap-frozen in liquid nitrogen and stored at −80°C until further analysis.

### 2.2 High-resolution HLA typing

Total DNA was extracted from 5 x 10^6^ cells using the DNeasy Blood & Tissue Kit (Qiagen) according to the manufacturer’s instructions. Samples were then sent to the Australian Red Cross Lifeblood for Next-Generation Sequencing (NGS)-based profiling of HLA class I and II genotypes.

### 2.3 RNA Sequencing

We have used RNA Seq data previously published (24). Briefly, RNA was extracted with Qiagen RNeasy columns according to manufacturer’s instructions including on column DNase treatment. All RNAseq samples were submitted for sequencing in triplicates. The library preparations and RNAseq were performed at Beijing Genomics Institute with a minimum of 20 million read counts per sample using paired end 150bp sequencing. Ribosomal RNA was depleted and polyA RNA were enriched from the total RNA samples. The library was then sequenced on a DNBSEQ G-400 platform. Data processing was automated using the RNAsik pipeline. Differential expression analysis was performed using the Degust platform with the limma-voom method (25, 26). A permissive threshold was applied to determine significant differential expression of genes in the knockout (KO) groups compared to the parental lines, defined log2 fold change >1 or <−1, with adj.p < 0.05.

### 2.4 Flow Cytometry

Cells cultured under basal conditions or treated with 5 ng/mL interferon-gamma (IFNγ) (Sino Biological) for 48 hours and were stained with a pan-HLA class I antibody (W6/32) conjugated to FITC (Leinco Technologies). Fluorescence intensity was assessed by flow cytometry to evaluate changes in cell surface HLA-I expression. All flow cytometric assays were acquired using a BD LSRFortessa™ cell analyser at the FlowCore facility, Monash University. Data were analysed using FlowJo® software version 10.10.0 (Becton, Dickinson and Company, Ashland, USA). A statistical analysis was conducted using GraphPad Prism software, version 10.4.1.

### 2.5 Purification of HLA-peptide complexes from cell pellets

HLA-peptide complexes were isolated using the semi-automated, SAPrlm, method on the KingFisher platform, as previously described (27). Briefly, cell pellets were lysed in 500 µL of lysis buffer containing 1% CHAPS (Thermo Scientific), 50 mM Tris (pH 8.0), 150 mM NaCl, 10 mM 2-chloroacetamide (Sigma Aldrich), and 1× Halt™ Protease and Phosphatase Inhibitor Single-Use Cocktail (Thermo Scientific). Lysates were gently mixed and incubated on a roller at 4°C for 1 h.

HLA-peptide complexes were immunoprecipitated using hyper-porous magnetic protein A beads (MagReSyn, ReSyn Biosciences) conjugated with the pan anti-HLA-I antibody W6/32 (Leinco Technologies). Peptide ligands were eluted under acidic conditions and subsequently desalted and concentrated using C18 resin columns (BioPureSPN) to remove buffer components, detergents, and salts that could interfere with downstream LC-MS/MS analysis.

To enhance the depth of peptide identification by LC-MS/MS, peptides were fractionated using stepwise elution with acetonitrile (ACN) at 5, 10, 15, 20, 25, and 28%. To maximise the recovery of distinct peptide species while maintaining sample complexity for optimal MS analysis, fractions were pooled as follows: 5% and 20% (Pool 1), 10% and 25% (Pool 2), and 15% and 28% (Pool 3). Fractions were then dried under a vacuum concentrator (Labconco, USA) and stored at -80°C until LC-MS/MS analysis to preserve sample integrity and prevent degradation.

### 2.6 LC-MS/MS Data Acquisition

Peptides were analysed using a Data-Dependent Acquisition (DDA) approach. Prior to LC-MS/MS, peptides were reconstituted in 2% acetonitrile (ACN)/0.1% trifluoroacetic acid (TFA) and loaded onto a trap-and-elute system consisting of a Thermo Fisher Scientific Acclaim™ PepMap™ 100 nanoViper C18 trap column (50 mm × 300 µm, 5 µm, 100 Å) and an Acclaim™ PepMap™ 100 C18 analytical column (50 cm × 75 µm, 2 µm, 100 Å) on an Orbitrap Exploris 480 mass spectrometer coupled to an UltiMate 3000 RSLCnano UHPLC system. Peptides were separated using a 90-minute linear gradient from 6% to 36% buffer B (80% ACN/0.1% formic acid), followed by an increase to 99% buffer B at 103 minutes, held for 5 minutes, at a flow rate of 250 nL/min. MS1 spectra were acquired in the m/z range of 350–1700 at a resolution of 120,000 with an RF lens setting of 40%. Monoisotopic peak determination was set to Peptide with relaxed restriction. The automatic gain control (AGC) target was set to 250% or until reaching a maximum injection time of 50 ms. Precursor ions with charge states +1 to +4 were selected, and dynamic exclusion was applied for 10 seconds within a 15-ppm mass tolerance. For MS2 scans, precursor ions were isolated using a 1.1 m/z window and fragmented by higher-energy collisional dissociation (HCD) at a normalized collision energy of 30%. Fragment ions were detected at a resolution of 15,000 with a scan range mode set to ‘define first mass’ at 110 m/z. The AGC target was set to 200% or until reaching a maximum injection time of 100 ms.

### 2.7 HLA Peptide Identification

MS/MS spectra were analysed using PEAKS 11 software (Bioinformatics Solution Inc). Raw MS files were imported with a parent mass error tolerance of 15 ppm and a fragment mass error tolerance of 0.02 Da. Protein grouping was disabled to allow all potential protein sources for a given peptide sequence to be annotated.

An initial de novo sequencing of MS/MS spectra was performed, followed by a database search against the human proteome. The search parameters included variable post-translational modifications (PTMs): oxidation of methionine (M), acetylation of lysine (K), carbamidomethylation of cysteine (C) as a fixed modification, and cysteinylation, deamidation (N/Q), and methylation (K/R) as variable modifications. A false discovery rate (FDR) threshold of 1% was applied at the peptide level, and all confidently identified peptides were exported for further analysis.

### 2.8 Peptide Binding Prediction Using NetMHC pan 4.1 and Peptide HLA Binding Modelling

Peptide binding predictions were performed using NetMHC pan 4.1 (28), a tool based on artificial neural networks that predicts immunopeptide binding affinity to HLA allele molecules. The default binding affinity cut-off of 2.0 was applied to classify peptides as pHLA binders or not assigned. AlphaFold3 was used to predict the structure, and PyMOL was utilized to examine the HLA-peptide interaction within the binding groove to further validate the peptide-HLA binding (29).

### 2.9 Criteria for selection peptides for immunogenicity assays

To qualify as pHLA antigens, peptides were required to map to a single source protein and be absent in the human benign atlas (30). As a final quality control step, only peptides uniquely identified in parental cells, derived from TAA and supported by high confidence MS2 spectra were selected for synthesis and spectral validation prior to immunogenicity assays. The TAA database, comprising 129 entries, was curated in-house based on an extensive literature review of reported genes associated with gliomagenesis and oncogenic pathways (Supplementary Table S1). Data sources included The Cancer Genome Atlas (TCGA), the cancer proteome in Human Protein Atlas (HPA) (31) and Childhood Cancer Model Atlas (CCMA)(24). The expression of gene sources for the selected peptides was screened on tissue samples to ensure adequate presentation levels in actual tumour tissue. For RNA isolation, fresh frozen DMG brainstem tumour tissue samples (5 mg) from 39 patients were processed. RNA was extracted by using the RNeasy Plus Micro Kit (Qiagen, 74034). Nanodrop and Qubit were used for RNA concentration measurement and a minimum of 200 ng RNA was used for the library preparation (RNA-seq - TruSeq mRNA-seq library). RNA-sequencing was done by Illumina NovaSeq 6000, 50PE sequencing to 25M clusters per library. RNA-seq data was obtained as FASTQ files from all samples and processed using the nf-core RNA-sequence pipeline (v 3.14.0) with default parameters (32). Gene expression quantification was performed using Salmon (version 1.4.0), resulting in a count matrix used for differential expression analysis using the DESeq2 package (v. 1.32.0) (33, 34).

### 2.10 Immunogenicity assay of selected peptides

Fresh donor peripheral mononucleated cells (PBMCs) were isolated from buffy coats using Ficoll-Paque density centrifugation and cryopreserved in 90% foetal calf serum (FCS) and 10% dimethylsuphoxide (DMSO) and stored in liquid nitrogen. For antigen-specific T cell expansion, 0.25x10^6^ PBMC per well were cultivated in 200 µL of AIM V medium in 96-well round bottomed plates in the presence of 1 µg/mL peptide. In specific experiments, unrelated immunogenic peptides (Flu NP HLA-A*01:01) were also used to generate specific T cell cultures as a control of T cell functionality. Recombinant human interleukin (IL)-2, IL-7 and IL-15 were added one day after. A quarter of cell culture media in each well was replaced with fresh AIM V media supplemented with IL-2, IL-7 and IL-15 every second day. Cultures were harvested on day 14 for functional assay. Specific cytotoxic activity of cultured T cells against autologous PHA T blasts un-pulsed or pulsed with relevant peptide was measured by standard ^51^Cr release assay (35). The percentage of specific lysis was calculated using the formula: (Chromium release for condition of interest – Chromium release in spontaneous release wells)/ (max chromium re-lease – Chromium release in spontaneous wells) x 100.

The same cytotoxicity assay was used to investigate the ability of peptide-specific T cells to lyse the parental and the H3K27M-KO cell lines. The natural killer cell-sensitive K562 cell line was also used as a target to assess the level of non-specific killing.

## 3. Results

### 3.1 H3K27M Mutation Induces Global Transcriptional Reprogramming

To elucidate the global transcriptional changes associated with H3K27M mutation, we investigated the gene expression profiles of the models. In total, 16,667 and 16,884 protein-coding genes were detected as transcribed in BT245 and SU-DIPG13 pairs, respectively (Supplementary Table S2). Following the knockout of the K27M point mutation, 19.39% of detected genes in BT245 and 16.58% in SU-DIPG13 showed a significant twofold or greater shift in expression levels (Figure 1A). Amongst the differentially expressed genes, 622 showed a significant downregulation in both models following H3K27M-KO (Figure 1B). Gene ontology enrichment analysis of these 622 genes using Metascape (36) revealed their association with neuronal processes, including trans-synaptic signalling, modulation of chemical synaptic transmission, and neuron projection development (Figure 1C). These pathways are essential for proper neuronal communication, differentiation, and circuit formation during brain development. The downregulation of these genes following H3K27M loss suggests a potential shift away from an active neuronal or synaptic state. Significant alterations were also observed in antigen processing and presentation machinery (APPM)- related genes (Figure 1D), although the patterns of these changes were inconsistent across the two models, reflecting context-dependent regulatory effects of the mutation.

**Figure 1.**
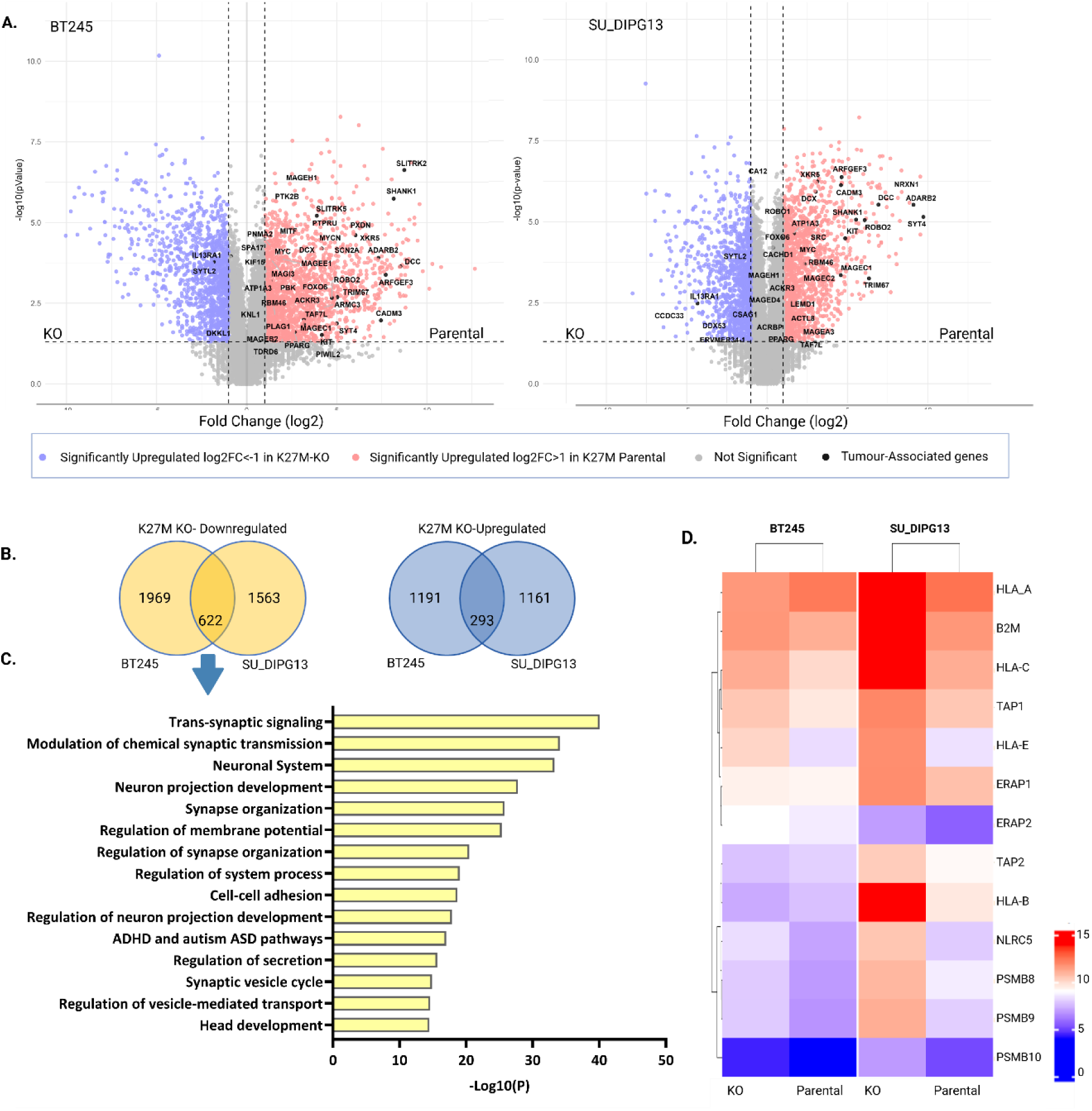
Transcriptomic analysis of isogenic parental (H3K27M+) and KO (H3K27M−) models of Diffuse Midline Glioma (DMG). A. Gene expression differentiation (log2 fold change) and statistical significance (-log10 p-value) in BT245 (left) and SU-DIPG13 (right), with highlighted Tumour-Associated Antigens (TAAs). B. Number of shared and unique differentially expressed genes between BT245 and DIPG13 parental and KO pairs (log2FC > 1 or < -1, p-value < 0.05). C. The top 15 enriched gene ontology terms for shared downregulated genes after H3K27M-KO. D. Differential expression of genes associated APPM, between H3K27M parental and KO for both cell lines.

### 3.2 H3K27M Mutation Does Not Directly Dysregulate HLA Class I Antigen Processing and Presentation Machinery

To determine whether the H3K27M mutation influences the HLA class I (HLA-I) APPM and alters the display of pHLA on DMG cell surfaces, we employed flow cytometry on paired model cell lines and assessed HLA-I expression using the pan HLA-I antibody W6/32. Initially, we assessed the basal HLA-I expression levels in the parental BT245 and SU- DIPG13 cell lines (Figure 2A and B, respectively). Flow cytometry analysis revealed that both cell lines were positive for cell surface HLA-I expression with no significant difference (p>0.05) between two cell lines (Figure 2C). However, when we compared the parental cell lines with their paired H3K27M-KO counterparts, we observed that H3K27M-KO increased HLA-I expression in SU-DIPG13 cells by ∼7-fold, whereas no change in HLA-I expression was observed in BT245 cells following H3K27M-KO (Figure 2C). We next investigated whether inflammatory cytokines enhance HLA-I expression in these cell lines and assessed whether the H3K27M mutation modulates the sensitivity of APPM to IFN-γ stimulation. Both parental BT245 and SU-DIPG13 cell lines showed a ∼3- and 4-fold increase in HLA-I expression, respectively, following 48 hours of treatment with 5 ng/ml IFN-γ (Figure 2D). Similarly, the H3K27M-KO cell lines showed HLA-I upregulation (∼3- and ∼2-fold, respectively) (Figure 2E).

**Figure 2.**
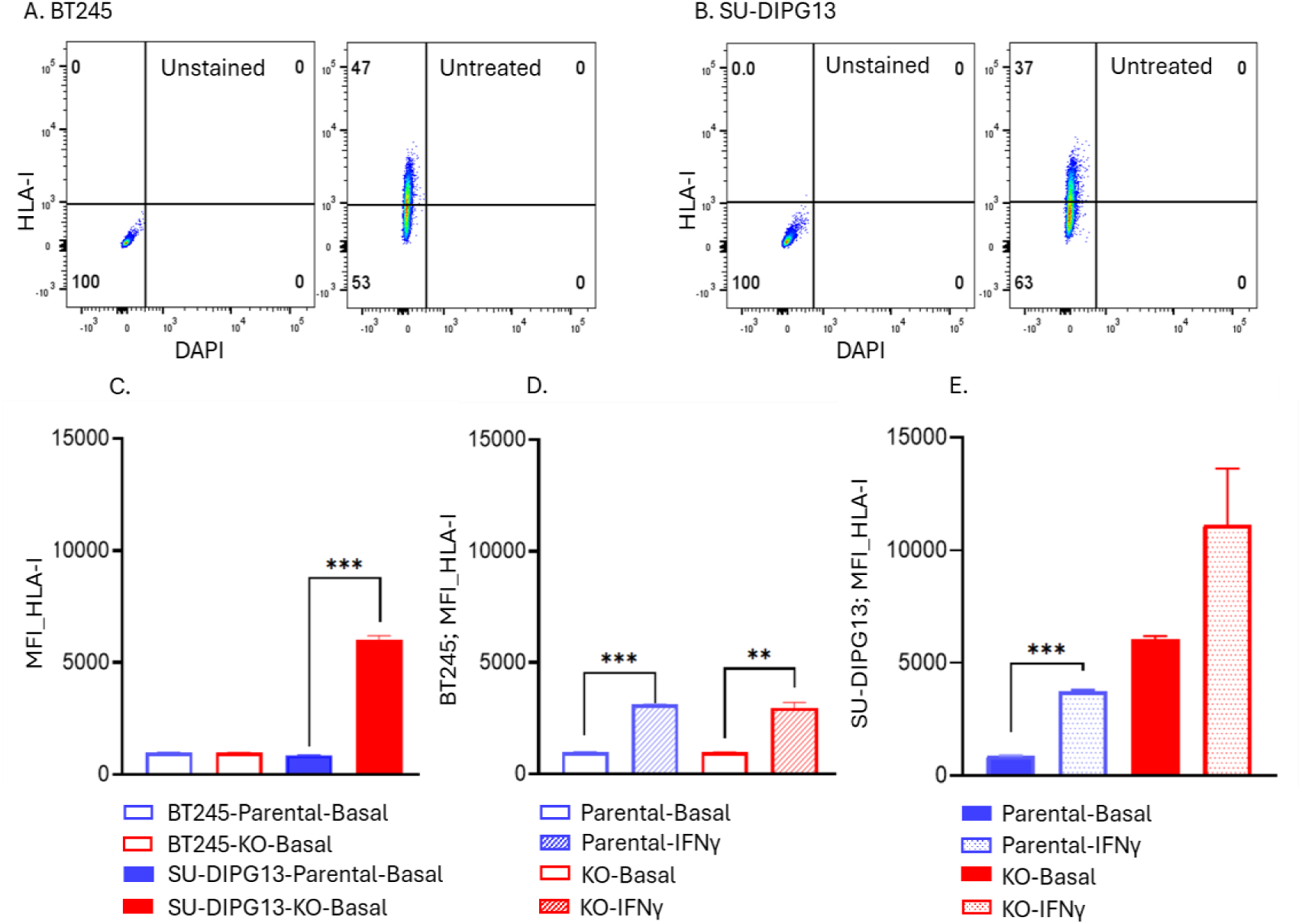
HLA-I expression in parental and H3K27M-KO DMG cell lines. (A, B) Representative flow cytometry dot plots for parental BT245 and SU-DIPG13 cell lines under unstained and basal conditions. The X-axis represents DAPI, indicating the live cell population, while the Y-axis represents HLA-I expression. The upper left quadrant indicates HLA-I-positive cells. The upper left quadrant indicates cells expressing HLA-I. Percentages reflect the proportion of cells within each quadrant. (C) Comparison of mean fluorescence intensity (MFI) of HLA-I expression in basal parental and H3K27M-KO BT245 and SU- DIPG13 cell lines. (D) BT245 parental and H3K27M-KO cells showed comparable upregulation of HLA- I expression following 48-hour treatment with 5 ng/ml IFN-γ. (E) SU- DIPG13 parental and H3K27M-KO cells also exhibited similar IFN-γ-induced upregulation of HLA-I expression (*p<0.05, **p<0.01, ***p<0.001).

### 3.3 The loss of H3K27M Modulates the DMG Tumour Antigenic Landscape

Although we did not observe a consistent relationship between the H3K27M mutation and HLA-I expression, our transcriptomic analysis revealed a higher expression of TAAs in H3K27M-mutant cell lines compared to their KO counterparts. Specifically, 24% and 17% of TAAs were significantly upregulated in H3K27M-mutant cells relative to their KO pairs (Figure 1A). Based on these findings, we sought to determine whether the H3K27M mutation influences the immunopeptidome repertoire of DMG cell lines by modulating TAA expression, thereby altering the antigenic landscape of these tumour cells.

To evaluate the presentation of pHLA derived from H3K27M-associated TAAs, we performed comprehensive immunopeptidomic analyses on parental and KO cell lines. Since IFN-γ treatment increase HLA-I expression across all cell lines, we also conducted immunopeptidomics analyses following IFN-γ stimulation. This approach allowed us to (a) increase the number of peptides identified from each cell line and (b) prevent misclassification of peptides as unique to a specific condition when, in fact, their low abundance under baseline conditions placed them below the detection limit of our mass spectrometry approach. Without IFN-γ treatment, such peptides might be missed, leading to an incomplete representation of the immunopeptidome. The overlap in pHLA repertoires between untreated and IFN-γ-treated conditions demonstrates that IFN-γ treatment expanded the pHLA repertoire. This expansion covers most peptides found in untreated conditions, especially in BT245, where the percentage of unique untreated peptides was minimal after stimulation (1.63% in parental and 1.59% in KO). In SU-DIPG13, IFN-γ had a less pronounced effect but increased the total number of peptides while maintaining a high proportion of shared peptides (63.03% in parental, 59.31% in KO) (Supplementary Figure S1). By combining peptides identified under both baseline and IFN-γ-treated conditions, we aimed to achieve a more comprehensive evaluation of the impact of the H3K27M mutation on the antigenic landscape of tumour cells.

In total, we identified 10,550, 10,107, 14,794, and 15,919 peptides from the BT245 parental, BT245 KO, SU-DIPG13 parental, and SU-DIPG13 KO cell lines, respectively (Supplementary Table S3-S6). Of these, 41.15% of peptides in the BT245 pair and 51.84% in the SU-DIPG13 pair were shared between the parental and KO cell lines. Consistent with flow cytometry data, the BT245 parental cell line displayed a comparable or greater number of identified peptides than its KO counterpart, whereas the SU-DIPG13 KO cell line exhibited a higher number of peptides relative to its parental line (Figure 3A).

**Figure 3.**
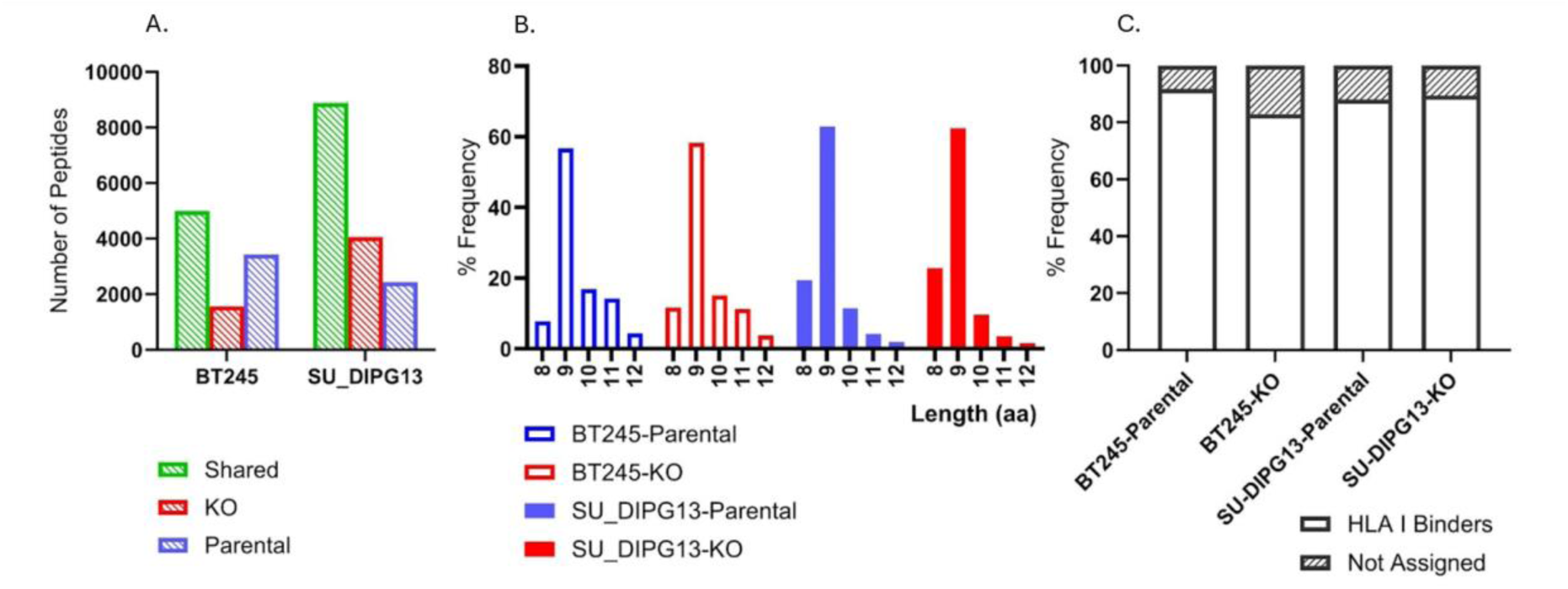
Comparative analysis of HLA-I peptides in BT245 and SU-DIPG13 parental and KO cell lines. (A) Number of HLA-I peptides identified in BT245 and SU-DIPG13 parental and KO cell lines, classified as parental-specific, KO-specific, or shared peptides. (B) Length distribution of HLA-I peptides in BT245 and SU-DIPG13. (C) Proportion of peptides assigned as binder to at least one HLA-I allele in BT245 and SU-DIPG13 cell lines.

More than 80% of the identified peptides in all samples fell within the canonical 8–12 amino acid range characteristic of HLA-I ligands (Figure 3B). Binding predictions using NetMHC pan 4.1, based on the HLA genotyping results (Supplementary Table S7), revealed that over 80% of peptides in each sample were predicted to bind at least one HLA-I allele expressed by the respective cell line (Figure 3C). These findings validate the robustness and accuracy of the immunopeptidomics data, demonstrating that the identified peptides align within expected HLA-I binding characteristics. Furthermore, we identified 5,584 unique protein sources in BT245 and 6,933 in SU-DIPG13, providing a broad landscape of antigen presentation across these models.

In the BT245 parental cell line, we identified 97 pHLA derived from 47 TAAs, accounting for 1.1% of the total immunopeptidome. In contrast, the KO counterpart exhibited 58 pHLA peptides from 28 TAAs, representing 0.8% of the immunopeptidome. Similarly, in SU- DIPG13, the parental line contained 133 pHLA from 53 TAAs, comprising 1.1% of the total immunopeptidome, whereas the KO variant displayed 137 pHLA from 45 TAAs, corresponding to 1.0% of the immunopeptidome. We further investigated whether specific TAAs exclusively contributed peptides in the parental cell lines but were absent in the KO counterparts, likely due to reduced or lost RNA expression of the corresponding TAAs (Figure 1A). In the BT245 pair, 22 TAAs were the source of 34 peptides detected only in the parental line (Figure 4A), whereas in SU-DIPG13, 14 TAAs exclusively presented 20 peptides in the parental cells (Figure 4B). Notably, seven peptides derived from shared TAAs were identified in both BT245 and SU-DIPG13 and were restricted to HLA-A*01:01. These findings suggest that the H3K27M mutation regulates the presentation of specific TAAs, leading to differential peptide presentation and shaping the immunopeptidome repertoire in an HLA allele-dependent manner.

**Figure 4.**
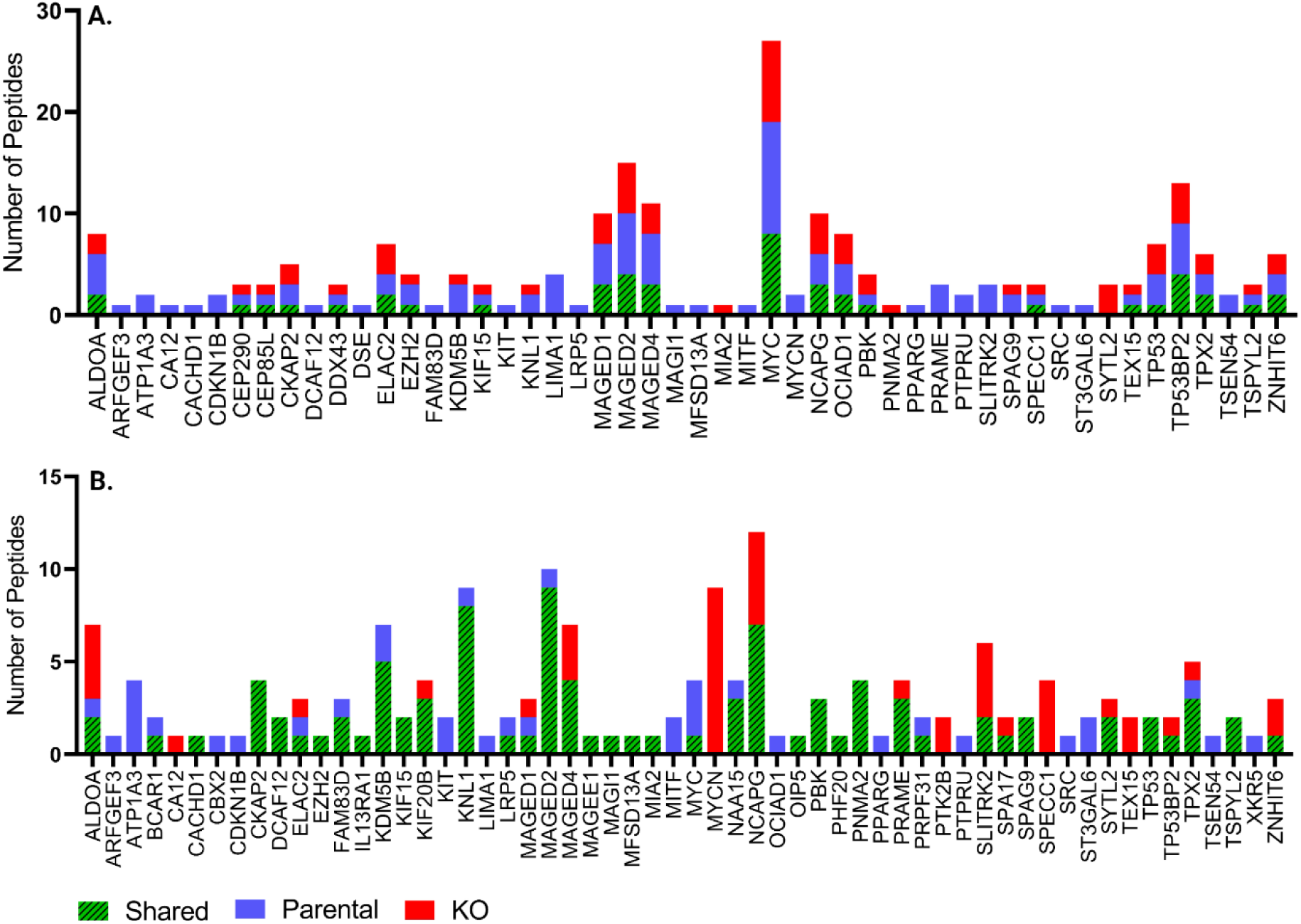
Contribution of Tumour-Associated Antigens (TAAs) to the antigenic landscape in (A) BT245 and (B) SU-DIPG13 parental (H3K27M+) and KO (H3K27M−) models. Bar graph showing the total number of HLA-I peptides derived from selected TAAs. Peptides are categorised as parental-specific (blue), KO-specific (red), and shared (green).

To identify candidate peptides for further validation and functional assays, we applied elaborate selection criteria to the 54 candidate peptides, prioritising a subset of 10 peptides based on their immunogenic potential. These selected peptides spanned five HLA types and included six peptides from SU-DIPG13 and eight from BT245, with four peptides shared between the two models (Table 1).

**Table 1.**
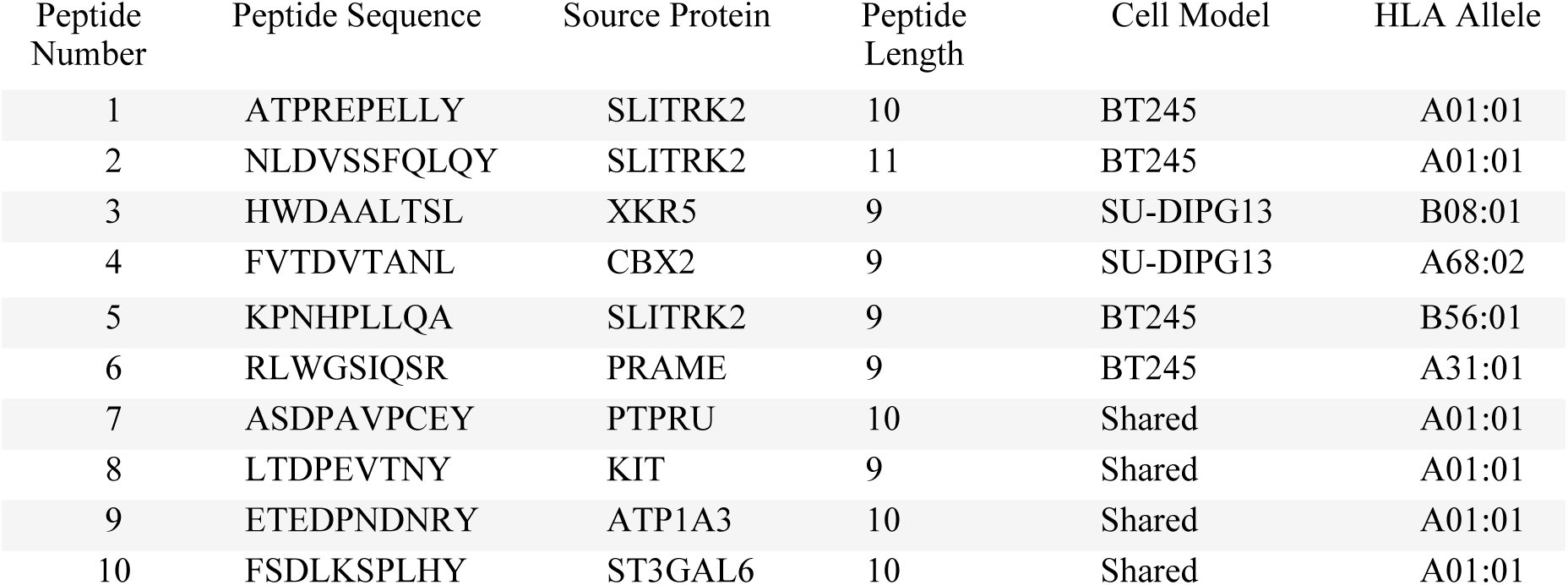
Peptides prioritized for validation and immunogenicity testing, selected based on exclusivity to parental cells, association with Tumour-associated antigens (TAAs), binding predictions to expressed HLA alleles, and higher RNA expression in H3K27M-DMG compared to non-malignant cells. Details include source proteins, associated cell models, HLA types, and peptide lengths.

First, to confirm the authenticity of these peptide sequences, we synthesized the selected peptides and performed MS2 spectral comparisons between the eluted peptides and their synthetic counterparts, ensuring sequence fidelity (Supplementary Figure S2). Next, we assessed their binding potential using in silico predictions, which confirmed that each peptide was a predicted binder to at least one HLA allele expressed in the respective cell lines. To ensure structural feasibility, we modelled the predicted peptide-HLA binding affinity using AlphaFold, which demonstrated high-confidence interactions within the HLA groove for all 10 selected peptides (Figure 5A and Supplementary Figures S3 A-J). Additionally, comparison with publicly available immunopeptidomics datasets confirmed that these peptide sequences were absent from previously reported HLA-bound peptides derived from benign tissues (30). Transcriptomic analysis revealed that the median expression of the source TAAs was significantly higher in a panel of H3K27M-DMG cell lines compared to non-malignant controls, as provided in the CCMA cohort (24). This suggests their tumour specificity and therapeutic relevance. Additionally, gene expression analysis of 38 DMG autopsy tissue samples confirmed the high expression of genes encoding the protein sources of the selected peptides, further validating their clinical significance (Supplementary Figure 4).

**Figure 5.**
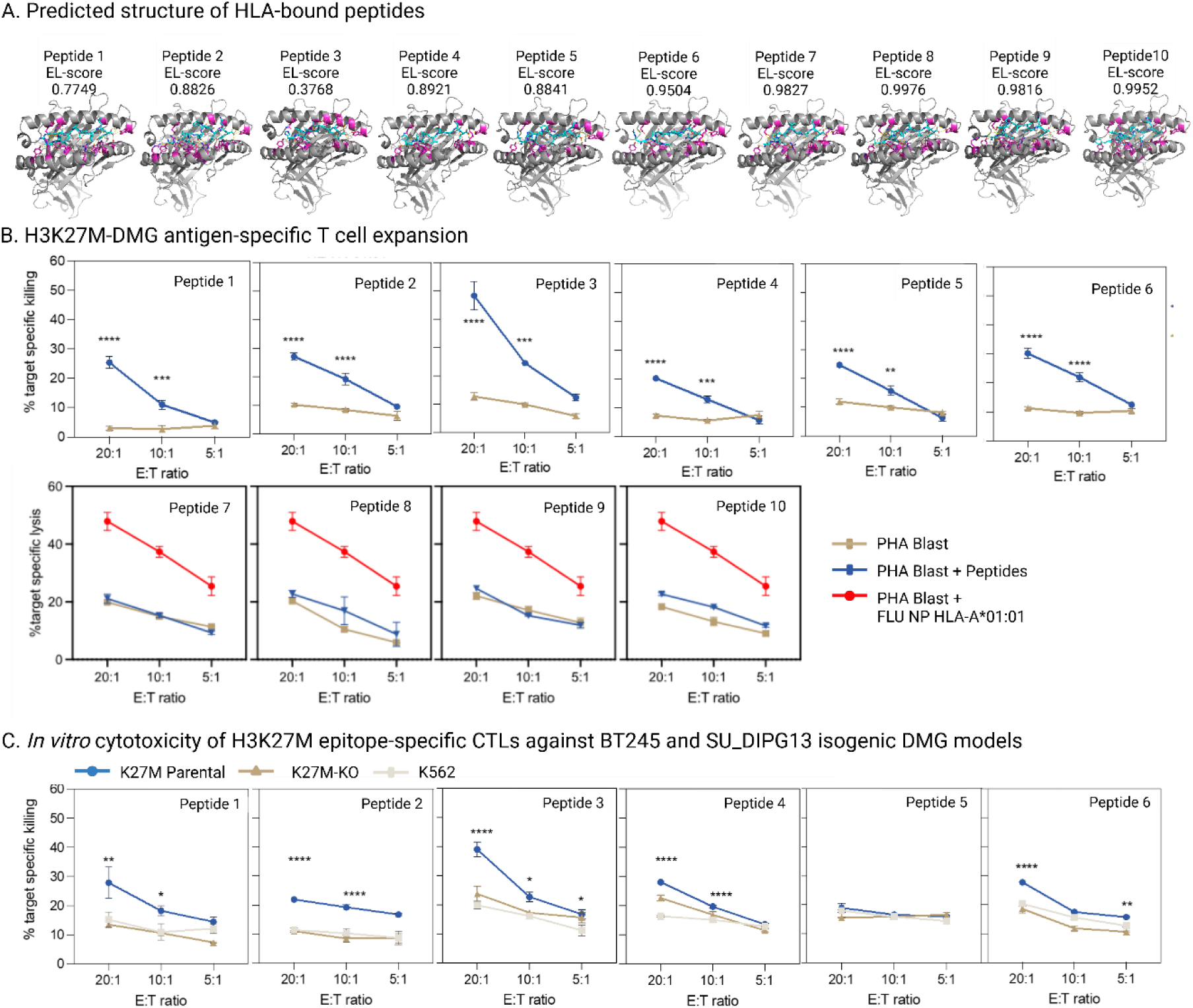
A) Structural representation of the HLA molecule bound to the peptide, predicted using AlphaFold3 with corresponding EL scores based on NetMHC pan 4.1, demonstrating their binding potential. The HLA heavy chain is shown in grey, with regions of predicted structural confidence highlighted in magenta. The peptide is depicted in cyan, positioned within the antigen-binding groove of HLA. Key interactions, including hydrogen bonds and molecular contacts, are indicated by yellow dashed lines. The model provides insights into peptide binding orientation and potential interactions critical for antigen presentation and T- cell recognition. B) Functional validation of the immunogenicity of peptides derived from H3K27M-DMG and identified by immunopeptidome. Peripheral blood mononucleated cells from 3 unrelated HLA-matched healthy donors were cultured for 14 days in the presence of each peptide. Expanded T cells were then assessed for their ability to lyse autologous PHA T blasts loaded or not with the corresponding peptides at the indicated effector:target ratios. For the 4 non-immunogenic HLA-A*01:01 peptides (Pep 7-Pep 10), the graphs also report the generation of T cells specific for the Flu NP HLA-A*01:01 epitope to ensure about the functionality of T cells. C) In vitro cytotoxicity of T cells specific for the 6 immunogenic peptides (Pep 1-Pep 6) against BT245 or SU_DIPG-13 cells. The cytotoxic responses of the same T cell cultures against K562 cells are also included as control for non-specific killing. Statistics: two-way Anova, Fisher’s LSD test for multiple comparisons vs negative controls; *p<0.05; **p<0.01; ***p<0.001; ****p<0.0001.

### 3.4 H3K27M Drives Presentation of Cytotoxic Epitopes from Specific Antigens

As a therapeutically relevant readout, peptide immunogenicity was functionally assessed by investigating their ability to elicit strong and specific cytotoxic T cell responses *in vitro*. To this end, we used viably frozen PBMCs from 9 healthy donors expressing HLA types (HLA-A*01:01, A*31:01, B*08:01, A*68:02, and B*56:01) matching those of the peptides identified in DMG cells. PBMCs from 3 HLA-matched healthy donors/peptides were cultured in the presence of each of the 10 selected DMG peptides for 2 weeks. T cells were then investigated for their ability to specifically lyse autologous PHA T blasts unpulsed or pulsed with each of the relevant peptides using standard cytotoxicity assays. As shown in Figure 5B, 6 peptides (peptides 1–6) were immunogenic, as T cell cultures demonstrated the ability to specifically kill peptide-pulsed PHA T blasts with a significantly higher specific lysis at 20:1 and 10:1 effector:target ratios as compared to unpulsed targets in all cases. These results were consistent among the T cell cultures generated from the three donors investigated, as shown by the limited inter-donor variability observed. In contrast, 4 peptides (peptides 7–10) failed to elicit T cells able to specifically kill peptide pulsed autologous PHA blasts, again with similar results obtained with PBMCs from the three donors tested. The ability of PBMCs of the same donors to generate T cells specific for an unrelated HLA-matched immunogenic peptide (HLA-A*01:01 Flu NP epitope) ensured the functionality of these cells and confirmed the lack of immunogenicity for peptides 7-10.

T cells specific for the 6 immunogenic peptides were also investigated for their ability to specifically lyse parental and H3K27M-KO DMG cell lines. As shown in Figure 5C, with the exception of T cells specific for peptide 5, all T cell cultures exerted significantly higher specific cytotoxicity against the parental as compared to the corresponding H3K27M-KO DMG cell lines. Notably, in all cases, the extent of killing of the H3K27M-KO cell lines was similar to that observed against the K562 cells included as control of non-specific lysis. These findings indicate that T cells specific for 5 of the 6 immunogenic peptides identified in the parental cell lines fail to recognise and kill the corresponding H3K27M-KO cell lines, consistently with the lack of detection of these peptides in the immunopeptidome of these cells.

These results demonstrate that the identified immunogenic peptides elicit T-cell responses capable of specifically recognizing and lysing H3K27M-mutant DMG cells. The absence of significant recognition in KO cell lines confirms that the H3K27M mutation is critical for the presentation of these neoantigens, underscoring their potential as targets for neoantigen-based immunotherapy.

## 4. Discussion

DMG remains one of the most aggressive paediatric brain tumours, with no effective standard of care beyond palliative radiotherapy (1, 23, 37, 38). The failure of chemotherapy and targeted therapies underscores the urgent need for precision medicine approaches guided by a deeper understanding of tumour biology. While DMG was initially classified within the glioblastoma (GBM) spectrum, mounting evidence has established pDMG-H3K27-altered as a distinct entity, differing from adult GBMs in its clinical presentation, developmental origins, and tumour microenvironment (39, 40). The discovery of H3K27M as a recurrent driver mutation has been pivotal in understanding the biology of DMG, as it fundamentally reprograms the epigenetic landscape. H3K27M disrupts PRC2-mediated H3K27me3, resulting in widespread transcriptional dysregulation and stalled differentiation pathways, which collectively sustain tumour growth and impact antigen presentation (4, 6, 7, 11, 41, 42).

Given that these epigenetic changes extend to antigen processing, we hypothesised that H3K27M could reshape the tumour’s antigenic landscape, influencing immune recognition and susceptibility to immunotherapy. To address this, we leveraged pDMG-H3K27M-KO models to investigate the relationship between H3K27M-driven epigenetic dysregulation and antigen presentation, with the goal of identifying actionable immunotherapeutic targets.

Our findings indicate that H3K27M alters antigen processing and presentation in a context-dependent manner. While PRC2 repression via EZH2, EED, and SUZ12 inhibition has been shown to de-repress MHC-I antigen presentation genes (42, 43), our analyses in isogenic DMG models (H3K27M+ and H3K27M-KO variants of SU-DIPG13 and BT245) revealed a more complex regulatory network. Flow cytometry demonstrated that H3K27M knockout led to increased HLA-I expression in SU-DIPG13 cells, whereas no significant change was observed in BT245 cells, suggesting that antigen presentation is regulated by additional factors beyond PRC2 inhibition. Interestingly, both parental and KO cells retained intact IFN- γ inducibility, supporting the feasibility of cytokine-based immunotherapies to enhance antigen presentation in DMG. Immunopeptidomic profiling further highlighted variability between models, with BT245 parental cells presenting a higher number of peptides than their KO counterparts, while SU-DIPG13 showed the contrary. This supports the notion that H3K27M’s influence on antigen presentation is modulated by tumour-intrinsic factors, such as lineage specification, differentiation status, and chromatin regulatory mechanisms. These findings are consistent with previous reports showing that the effects of H3K27M and H3K27me3 distribution are highly dependent on cellular context and neurodevelopmental timing, further highlighting the complexity of its role in antigen presentation (7, 42, 44).

Our data reveal a significant increase in TAA gene expression in parental K27M models (Figures 1A and 1B), indicating that the H3K27M mutation directly drives the upregulation of potential immunogenic targets. Consistently, our immunopeptidomics analysis identified numerous pHLA complexes, including a substantial fraction of novel peptides absent from the Immune Epitope Database (IEDB) (45), expanding the catalogue of DMG-associated antigens. Of the identified pHLAs from BT245 model, 8,356 peptides were previously documented in IEDB, while 2,439 were novel, including 1,372 DMG-specific peptides found exclusively in parental cells. In the SU-DIPG13 model, we identified 4,146 novel sequences and 864 unique to parental cells, with a markedly higher proportion of TAA-derived peptides in parental cells relative to KO conditions (Figure 4).

We selected 10 candidate peptides for subsequent immunogenicity assays to distinguish therapeutically relevant targets from those that are merely presented. Although we identified four shared peptides across both models, derived from the source antigens: cKIT, PTPRU, ATP1A3, and ST3GAL6, these peptides failed to induce T cell responses. This lack of T cell activation may result from low intrinsic immunogenicity or insufficient TCR affinity, underscoring the necessity for functional validation when selecting TAAs for immunotherapy. Beyond the broad effects on antigen presentation, the study identified six immunogenic peptides from four previously uncharacterized TAAs in DMG: SLITRK2, PRAME, XKR5, and CBX2 (Table 1). Among the identified peptides, only one peptide (Peptide 4, derived from CBX2), had been previously documented in the IEDB as a naturally presented ligand associated with cervical cancer and acute lymphoblastic leukemia under HLA-I restriction (46, 47). These peptides were validated using functional T cell assays, demonstrating their ability to expand T cells from HLA-matched donor PBMCs and elicit peptide-specific cytotoxic responses. Of note, CBX2, a key component of the PRC1 complex, plays a crucial role in maintaining transcriptional repression through chromatin remodelling and histone modification. By binding to H3K27me3, it modulates gene expression and lineage differentiation (48–50). The identification of immunogenic CBX2-derived peptide in the DMG immunopeptidome underscores the direct role of epigenetic dysregulation and chromatin regulators in shaping the tumour antigenic landscape, highlighting their potential as novel targets for immunotherapy.

Additionally, PRAME, a well-characterised cancer-testis antigen is particularly promising, given its restricted expression in normal tissues and high prevalence in multiple cancers (51–53). Current efforts focus on PRAME-specific immunotherapies, such as TCR-engineered T cell therapy (TCR-T), which has shown promising clinical responses in other malignancies (54). Furthermore, previous studies have reported PRAME-derived pHLAs expression in distinct patient-derived DMG cell lines, highlighting its relevance as a target antigen in H3K27M-DMG (21).

Despite the insights gained from this study, several limitations should be acknowledged. Our findings are based on *in vitro* models, and while isogenic H3K27M-KO cell lines provide a controlled system to dissect the effects of H3K27M on antigen presentation, they do not fully recapitulate the complexity of the *in vivo* tumour microenvironment. The immune landscape of DMG is influenced by factors such as stromal interactions, local immunosuppression, and blood-brain barrier constraints (9, 12–15, 55), which may further modulate antigen processing and presentation in a physiological setting. Additionally, our immunopeptidomic analysis was conducted on two DMG cell lines, which, while representative, do not capture the full heterogeneity observed across patient-derived tumours. While we identified novel immunogenic peptides, their functional impact on tumour immune responses and therapeutic efficacy in patients remains to be explored. Future studies should prioritise comprehensive immunogenicity assessments, including tumour-infiltrating lymphocyte responses and *in vivo* tumour clearance assays, to determine their clinical applicability.

Taken together, our study establishes a link between H3K27M-driven transcriptional dysregulation and tumour antigenicity, highlighting the potential of immunopeptidomics as a tool for identifying actionable immunotherapeutic targets. By leveraging epigenetically dysregulated antigens, we demonstrate that H3K27M creates distinct antigenic vulnerabilities that can be targeted using CD8⁺ T cell-based immunotherapies. Future studies should further evaluate the therapeutic potential of H3K27M-associated antigens, particularly in the context of adoptive T cell therapies and multi-epitope cancer vaccines. These findings lay the foundation for precision immunotherapy approaches in DMG, offering new hope for treating this otherwise intractable disease.

## 5. Conclusions

In conclusion, our study provides new insights into the immunopeptidome of H3K27M- DMG, revealing how epigenetic dysregulation influences antigen presentation and immune recognition. By leveraging immunopeptidomics and functional immunology, we identified novel TAA-derived epitopes that are selectively expressed in H3K27M-mutant tumours and functionally validated their immunogenicity. Our findings highlight PRAME and other dysregulated antigens as promising targets for HLA-restricted T-cell therapies, underscoring the feasibility of precision immunotherapy approaches in DMG. While further validation in patient-derived and in vivo models is necessary, our study establishes a foundation for antigen-directed immunotherapy strategies in this otherwise treatment-refractory disease. Future efforts should focus on integrating multi-epitope targeting, immune checkpoint modulation, and cytokine-based adjuvants to enhance therapeutic efficacy. These findings pave the way for the next generation of personalised immunotherapies aimed at improving outcomes for patients with H3K27M-DMG

## Acknowledgement

This study used BPA-enabled (Bioplatforms Australia) / NCRIS-enabled (National Collaborative Research Infrastructure Strategy) infrastructure located at the Monash Proteomics and Metabolomics Platform. We also thank Dr. Nada Jabado for providing the H3K27M knockout DMG models used in this study.

## Ethics Statement

All autopsy samples were collected after written informed consent from the patient’s guardian, as approved by the Institutional Review Board of Children’s National Hospital study entitled “Molecular Analysis of Pediatric Cancers” (#Pro00001339) or collected under the CNH-IRB approved exempt protocol “Biomarker Identification In Pediatric Brain Tumors” (#Pro00000747) or Swiss Ethical approval BASEC-Nr 2019-00615.

Healthy donor peripheral mononucleated cells were obtained from buffy coats provided by Australian Red Cross under the Biological Resources Agreement Agreement no: 24-08VIC- 06 – September 29, 2024.

## Conflict of Interest

The authors declare no conflicts of interest.

## Funding

This project was supported by grant 2022/GNT2019729 awarded through the National Health and Medical Research Council (NHMRC) and grant NCRI000108 awarded through The Medical Research Future Fund (MRFF). PF was supported by the Victorian Department of Health and Human Services through the Victorian Cancer Agency. We acknowledge the Isabella and Marcus Foundation for supporting this research through the Kye Funch Scholarship and the Tour de Cure Grant for their financial contribution.

## Authorship Contributions

PF and RD conceptualised the study. Methodology was developed by TS, CX, PD, RM, RD, TL, GG, ET, and BZ. Investigation was carried out by TS, TL, GG, GH, FF, NS, JEC, and JH. RNA sequencing was performed by TS, CX, and PD, while gene expression studies in tissue samples were conducted by BK, JN, and IW. Immunogenicity and cytotoxicity assays were carried out by BZ, RM, and RD. Resources were provided by RF, PF, RD, and RS. Data analysis and curation were performed by TS, BK, and JN. TS and PF wrote the original draft, with all authors contributing to review and editing. PF and RD supervised the study. Funding was acquired by PF, RD, and RF.

## Data Availability

All additional datasets supporting the findings of this study are provided in the Supplementary Data Excel file. Further information is available from the corresponding authors upon reasonable request.

---

**Figure S1.**
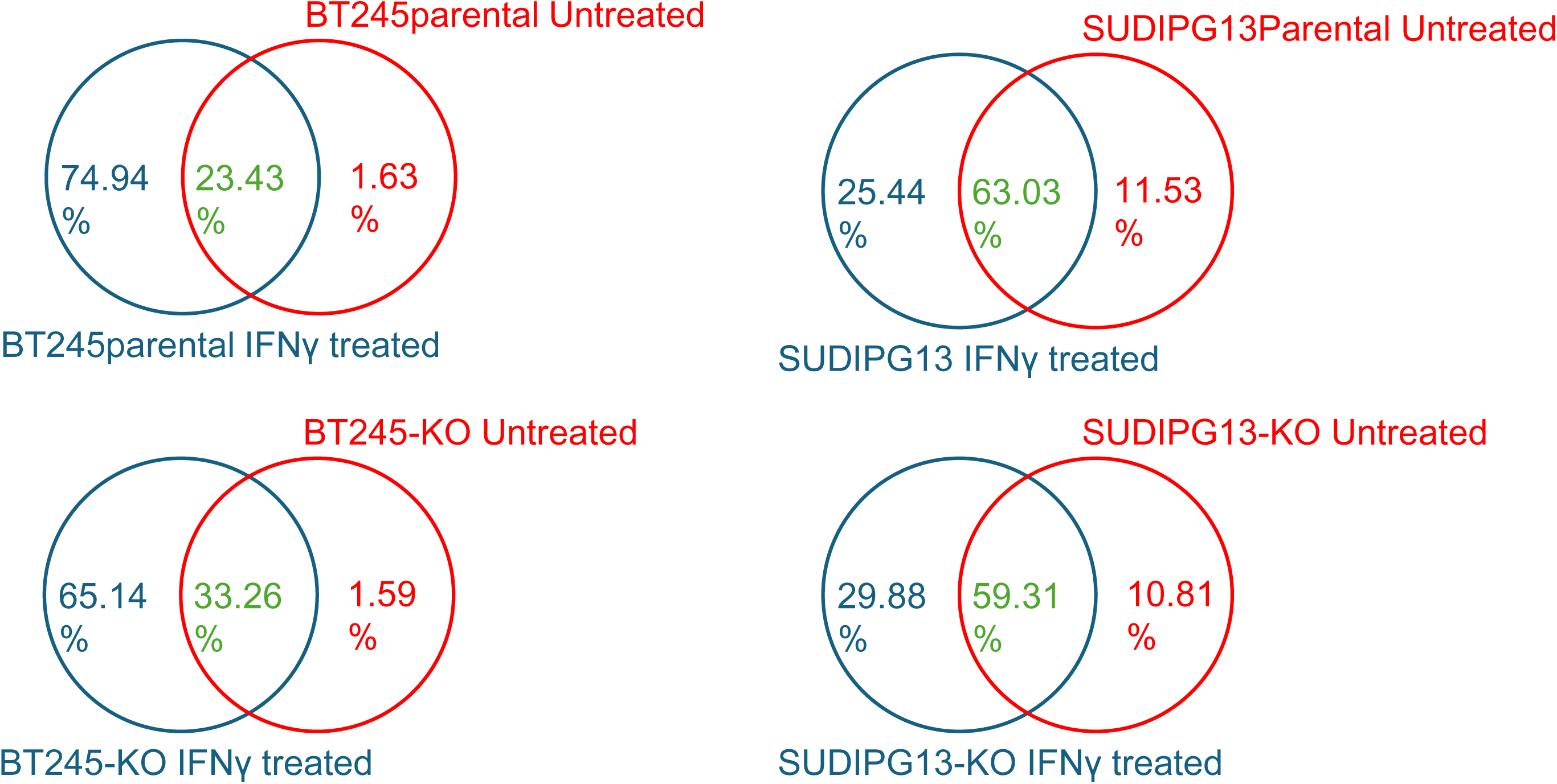
Comparison of pHLA Repertoires in Isogenic H3K27M-DMG Models Before and After IFN-λ Treatment The overlap of pHLA repertoires between untreated and IFN-λ-treated conditions in BT245 and SU-DIPG13 parental (H3K27M+) and knockout (H3K27M−) models. Percentages indicate the proportion of peptides unique to each condition (red and blue) and those shared between untreated and IFN-λ-treated samples (green). IFN-λ treatment significantly expanded the shared peptide repertoire, particularly in SU-DIPG13, while in BT245, it covered nearly all peptides present in the untreated condition. This highlights the impact of IFN-λ in enhancing antigen presentation.

**Figure S2.**
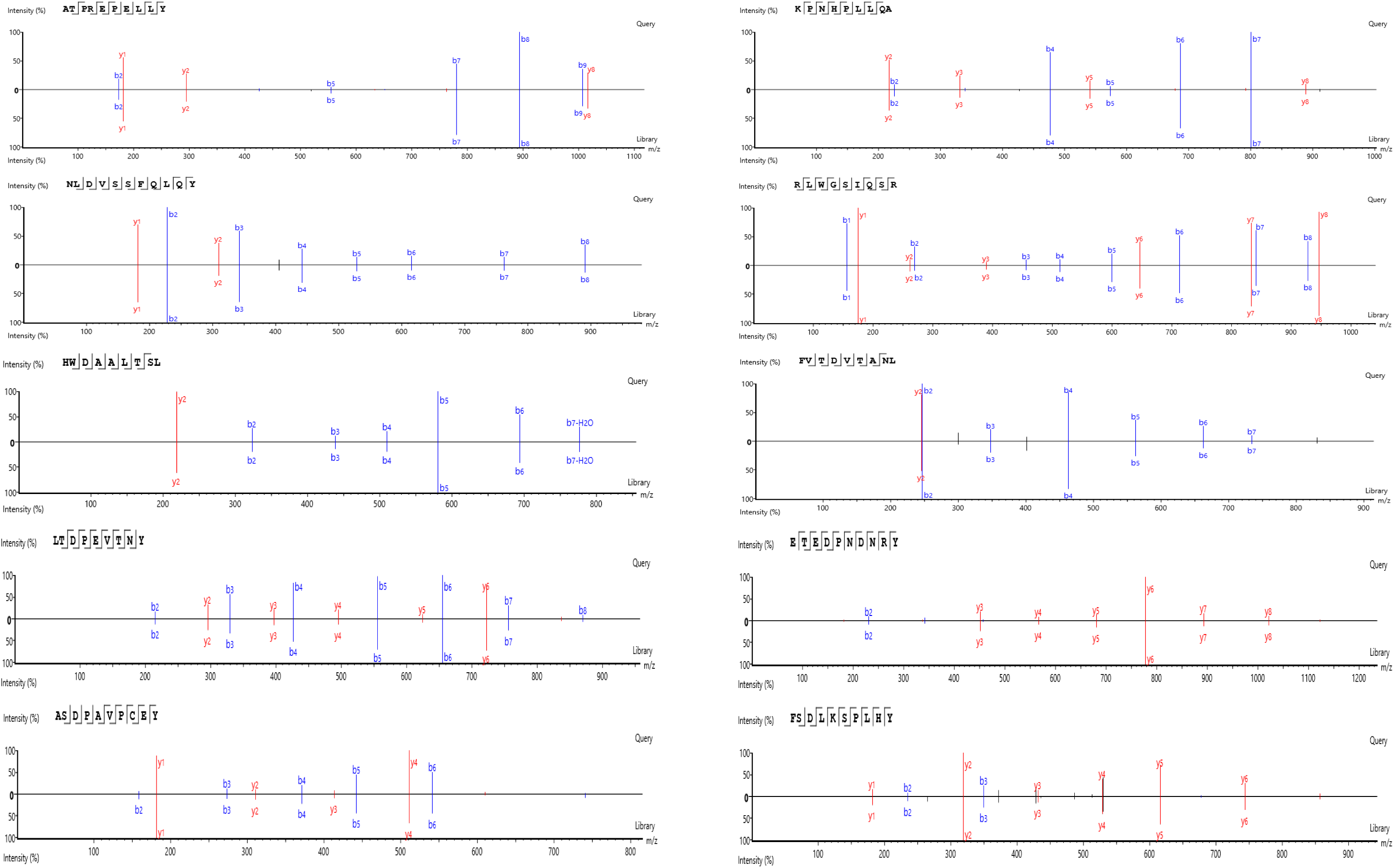
MS Spectra Validation of selected peptides. The actual spectra are shown on the top, while the synthetic spectra are displayed at the bottom (PEAKS 11 software (Bioinformatics Solution Inc).

**Figure S3.**
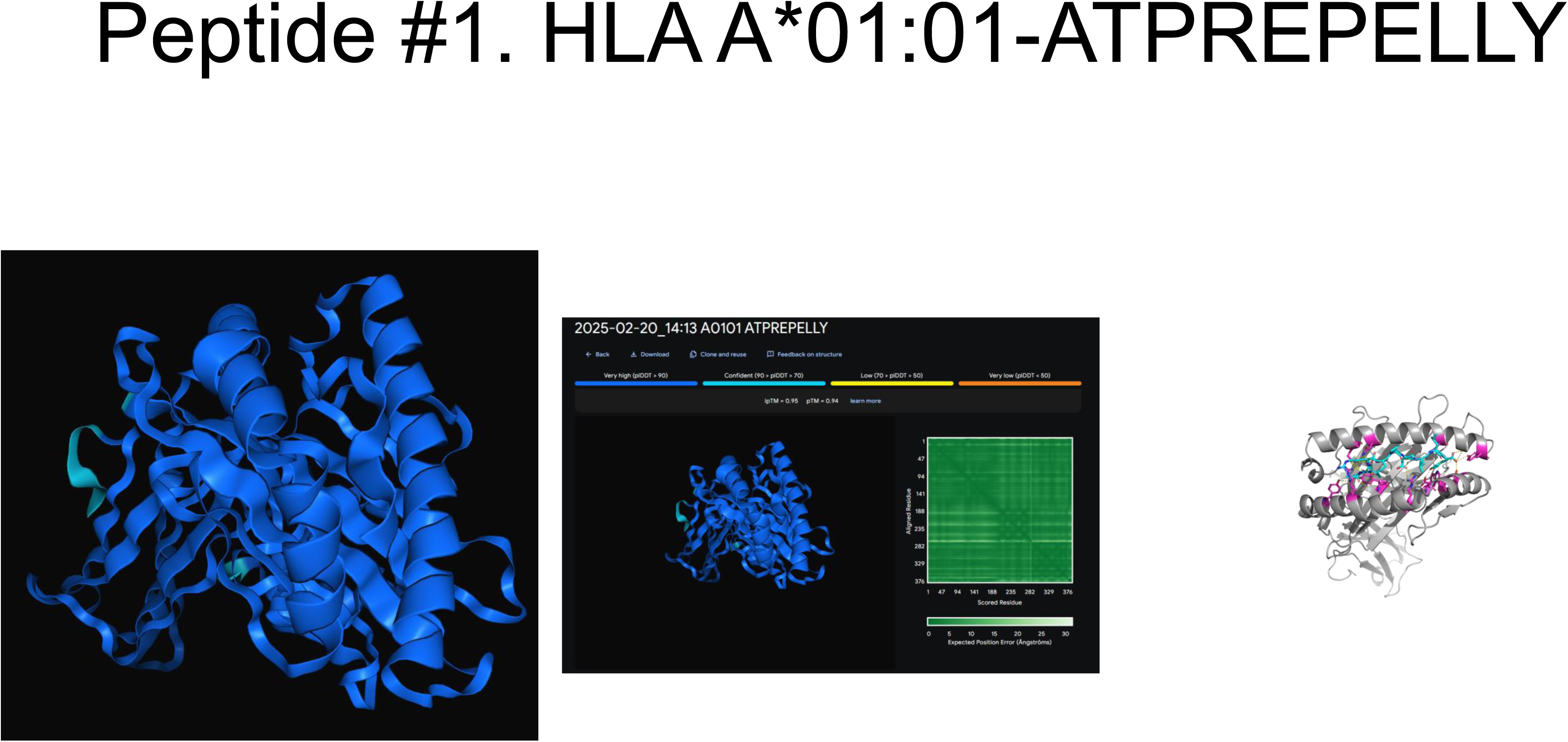
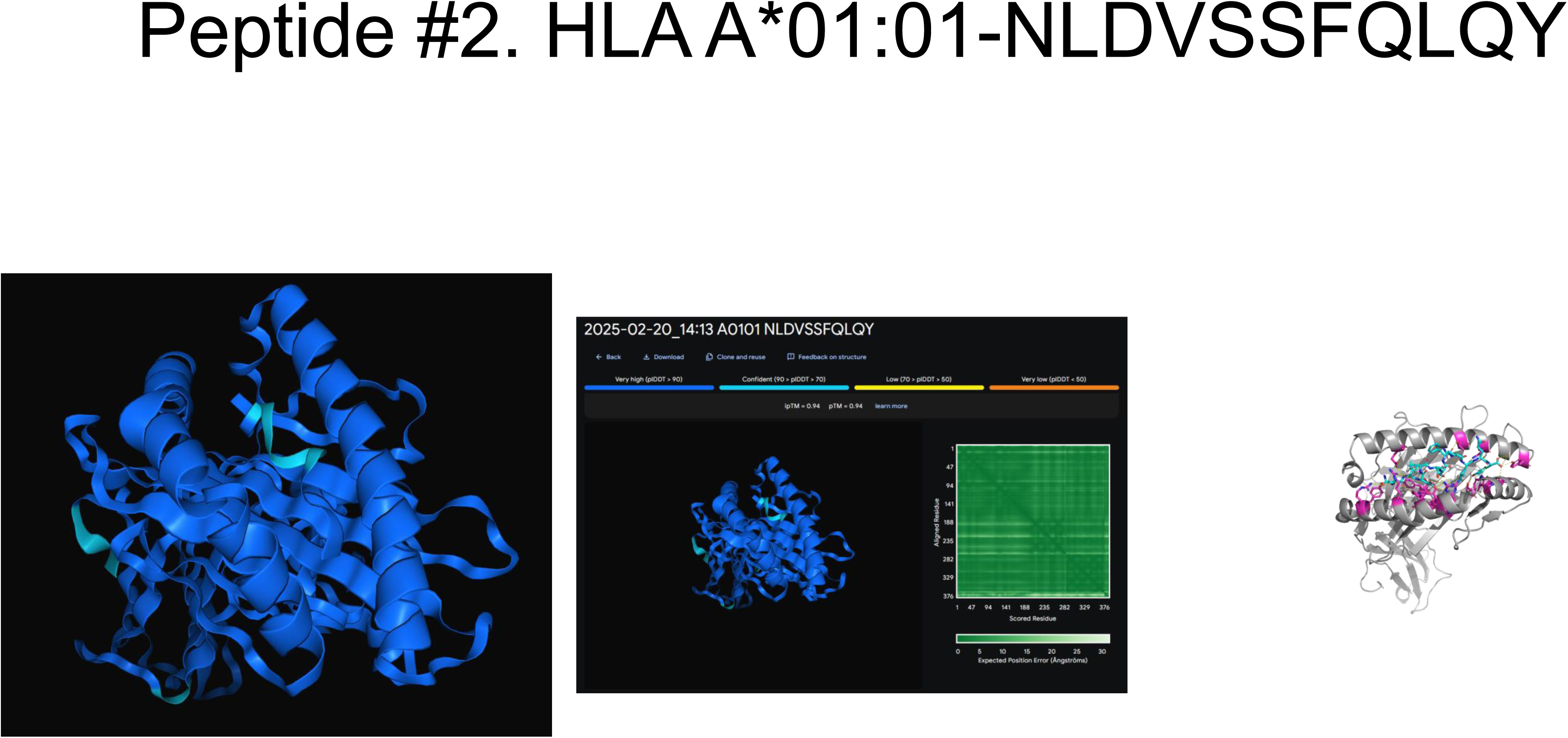
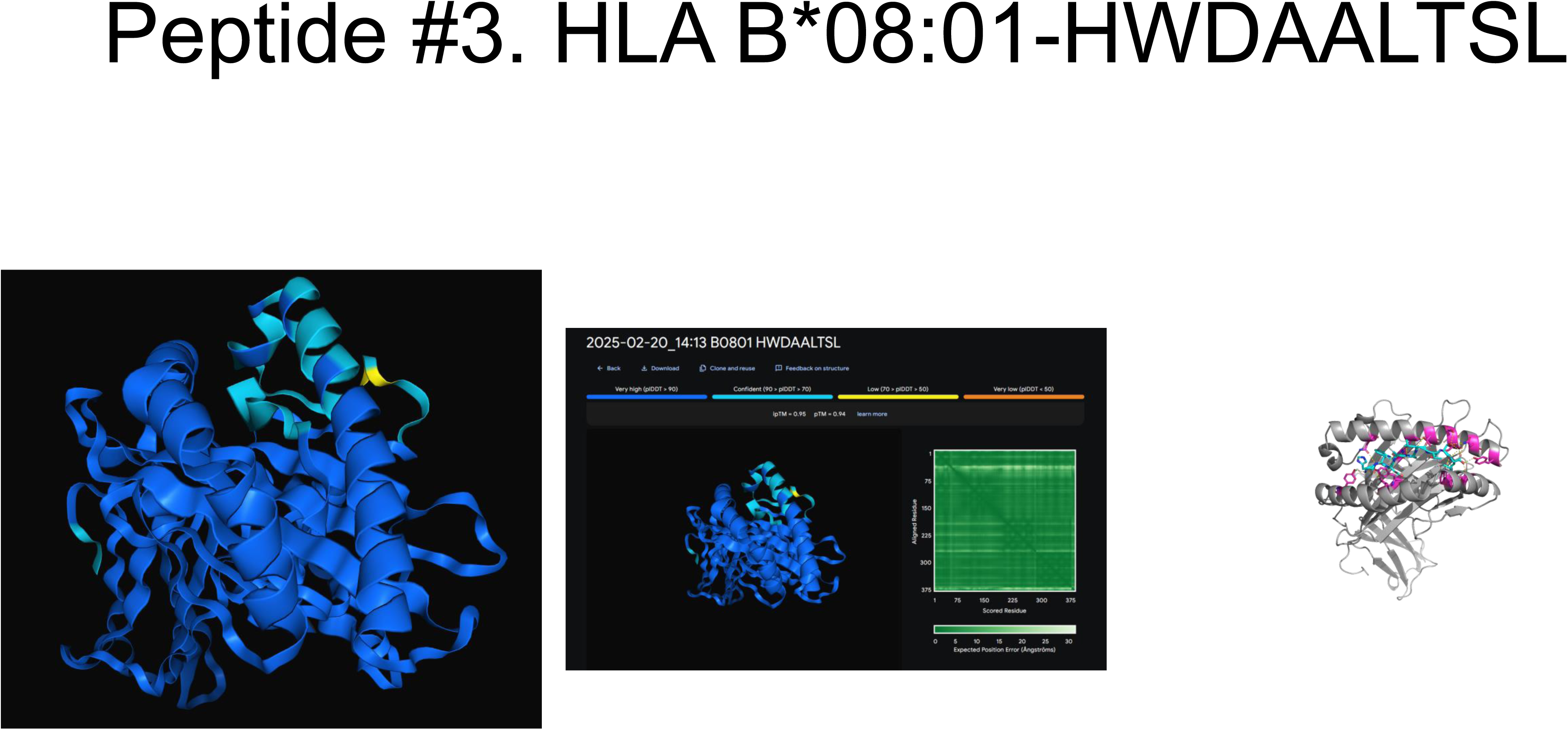
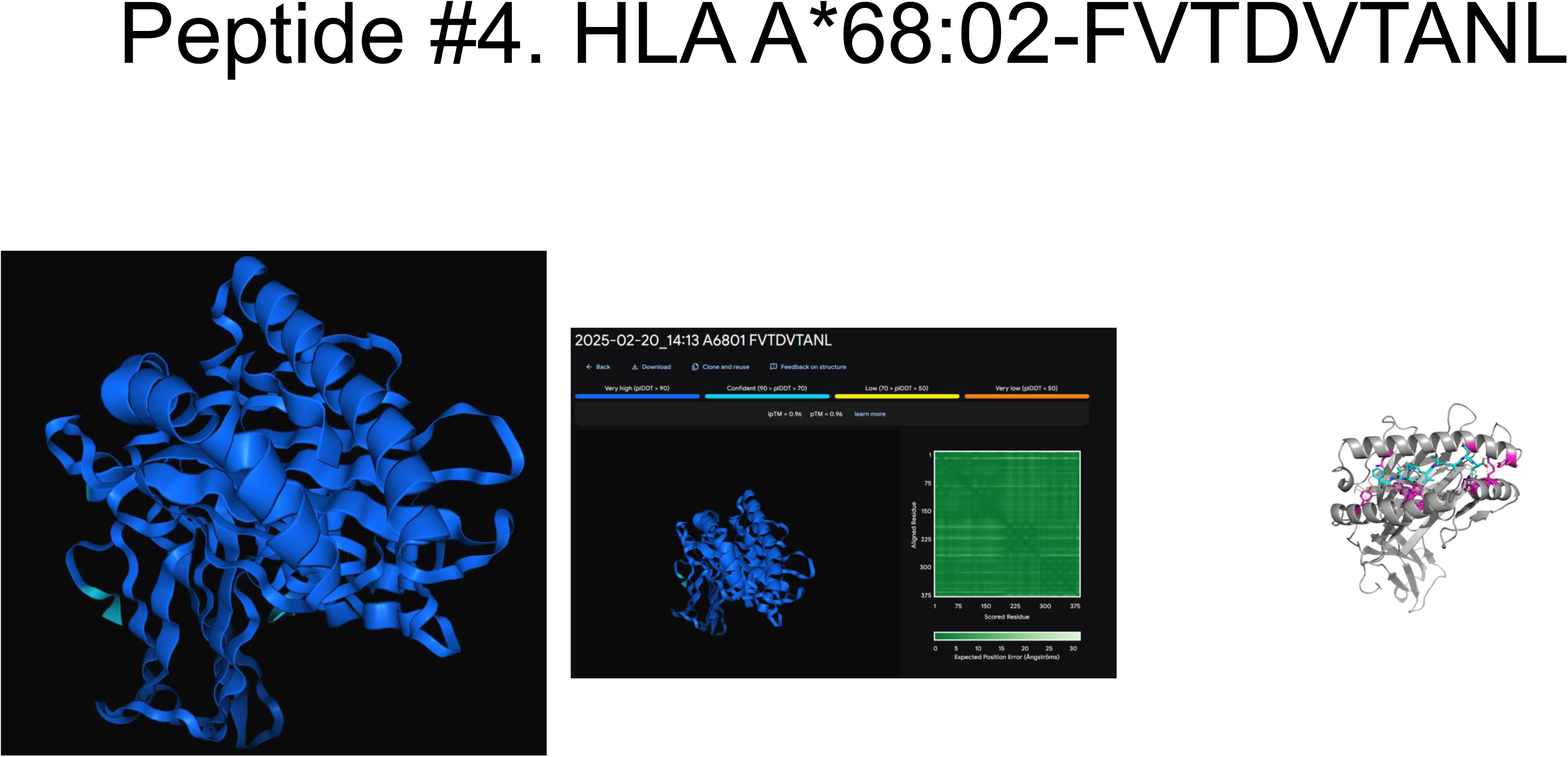
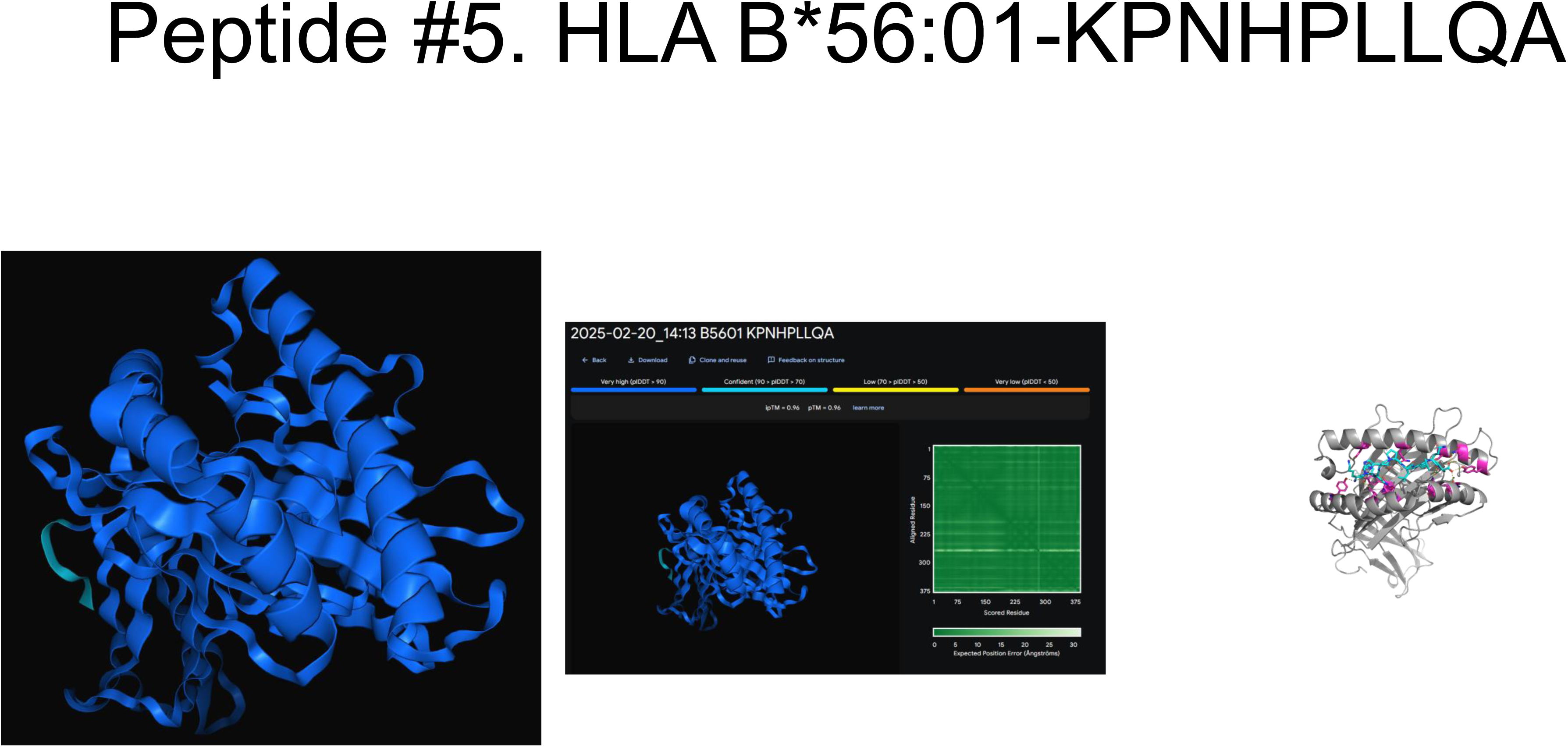
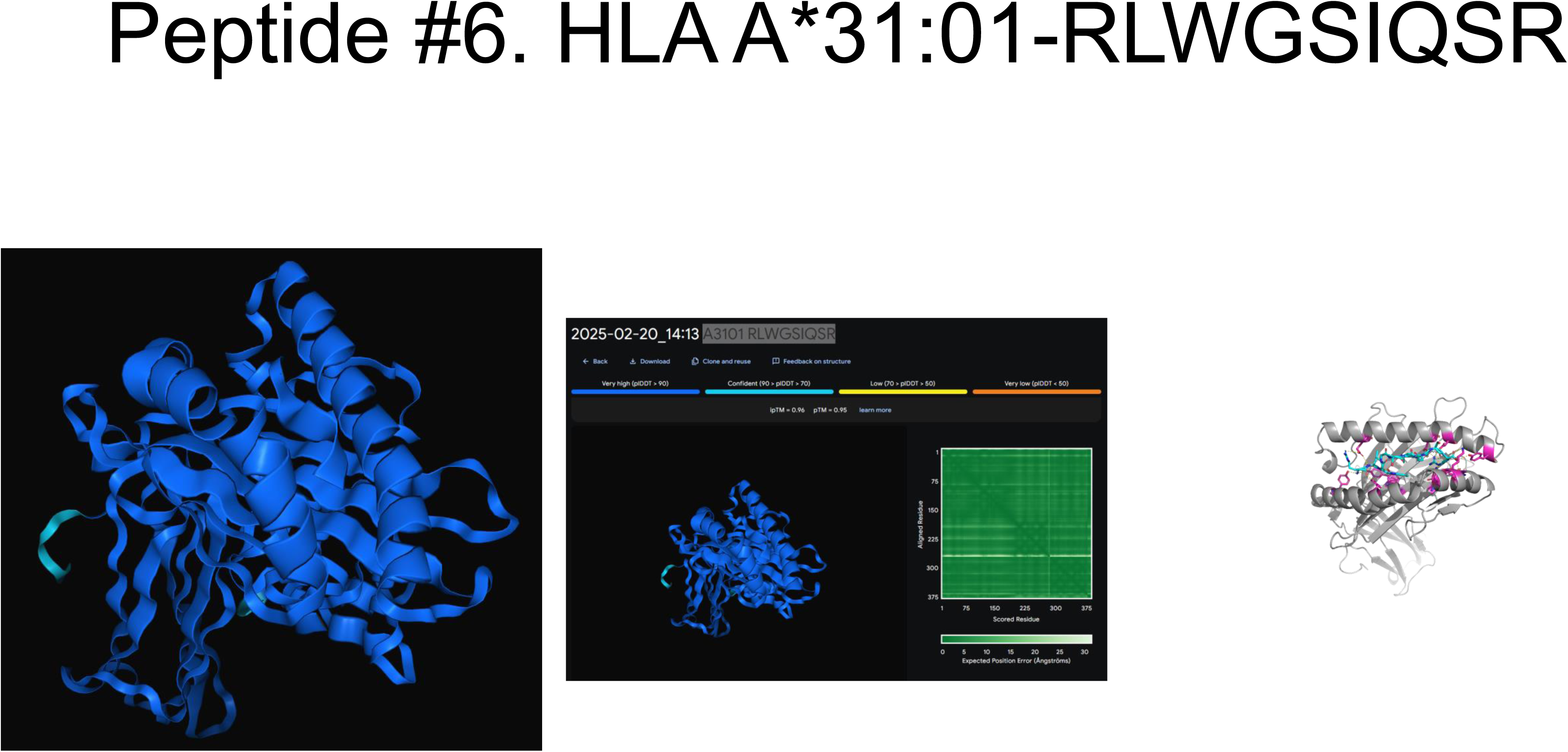
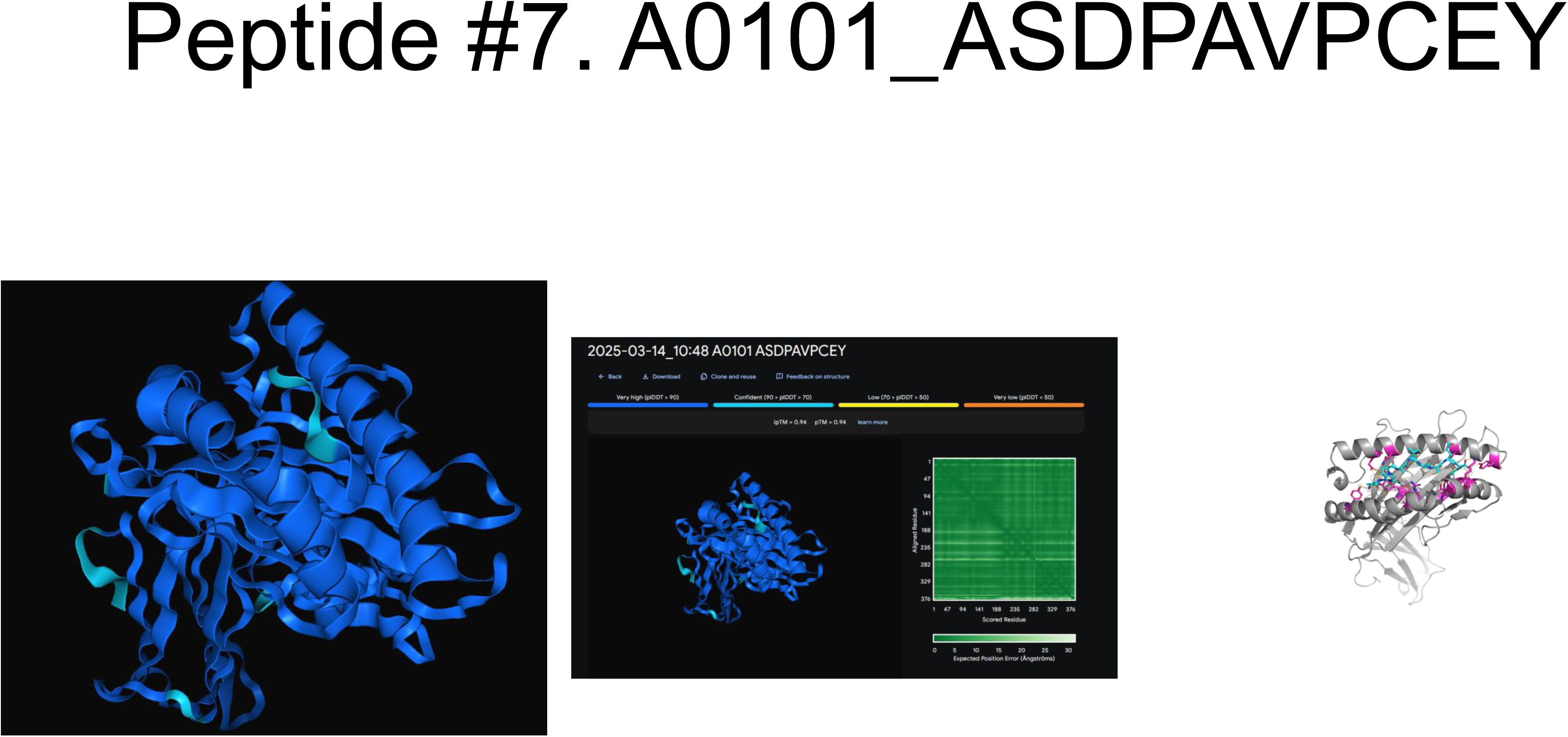
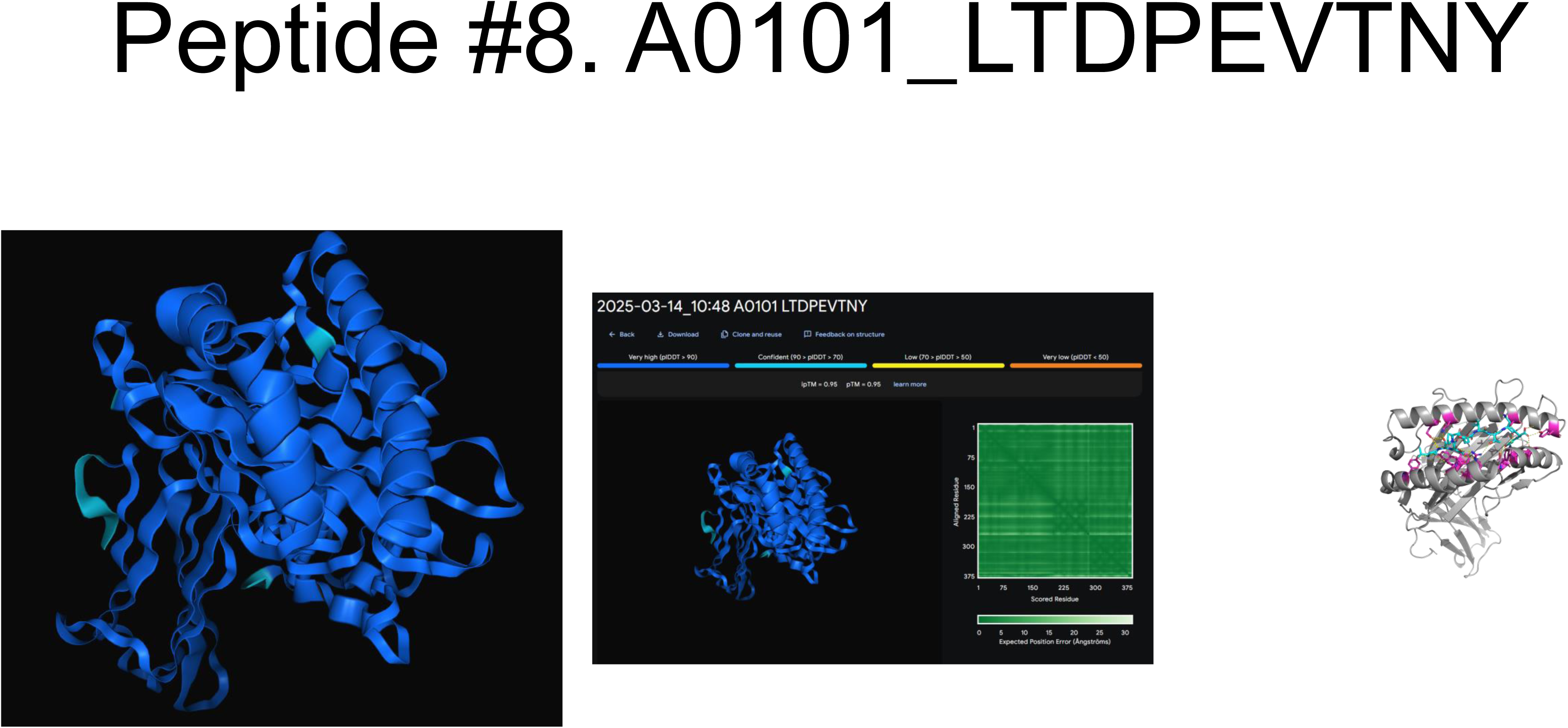
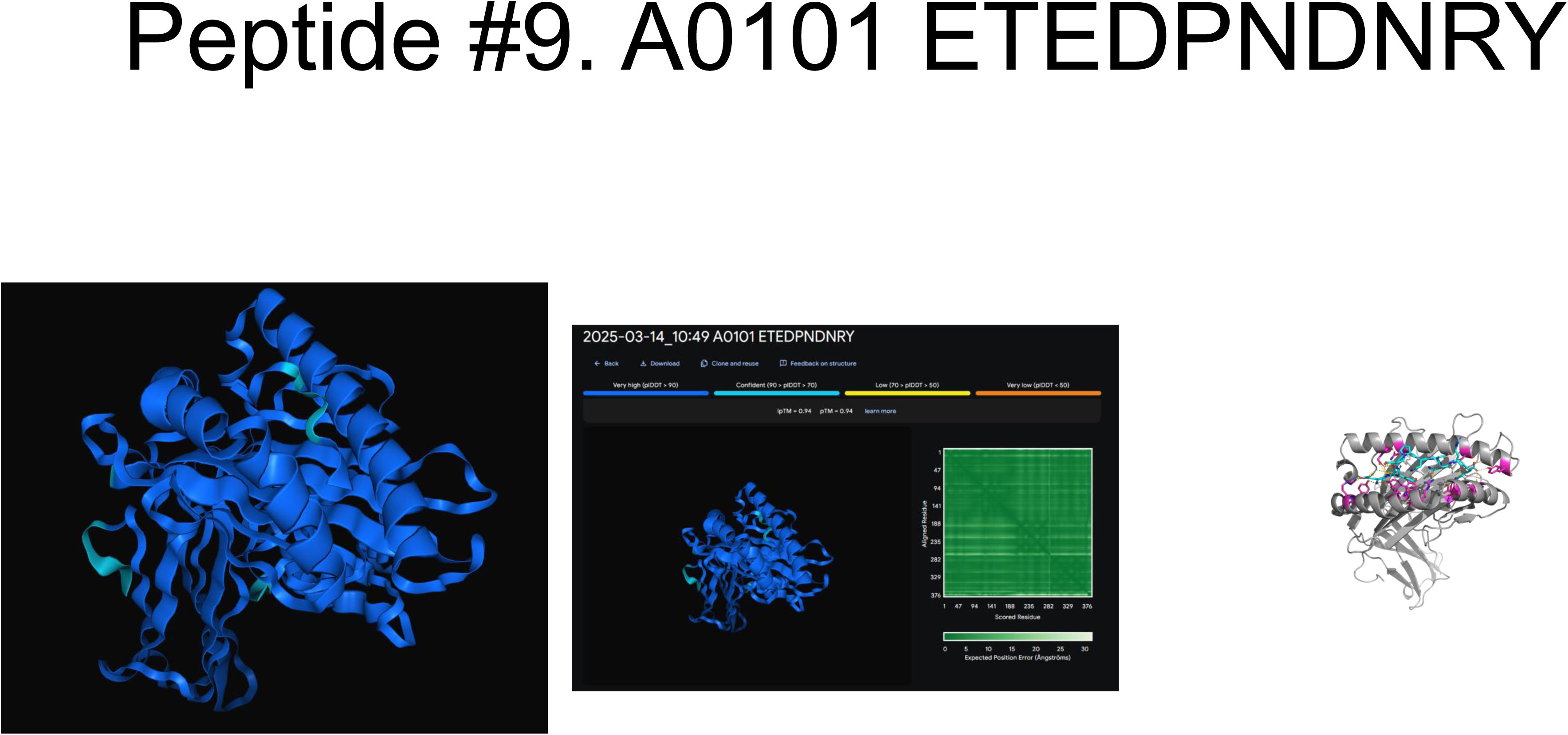
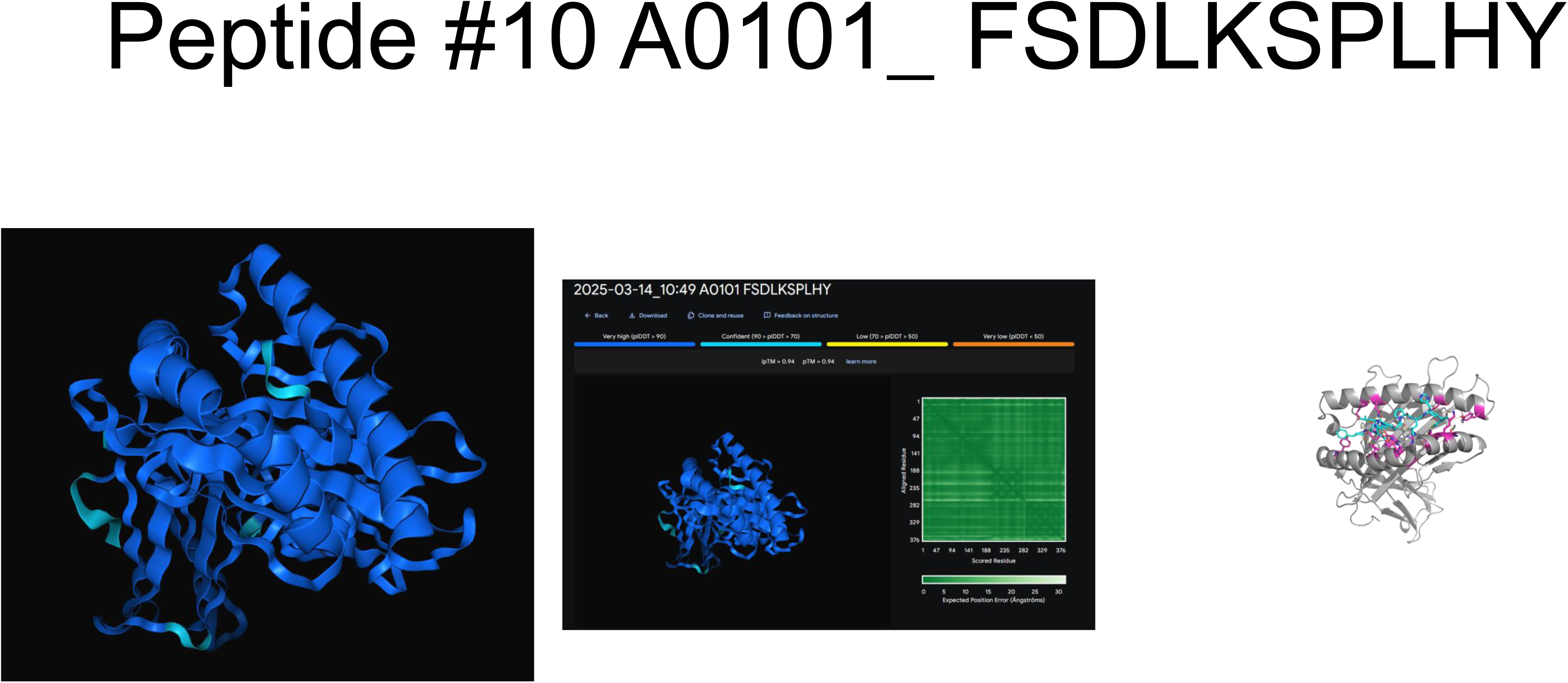
The PyMOLMolecular Graphics System, Version 3.0 Schrödinger, LLC.

**Figure S4.**
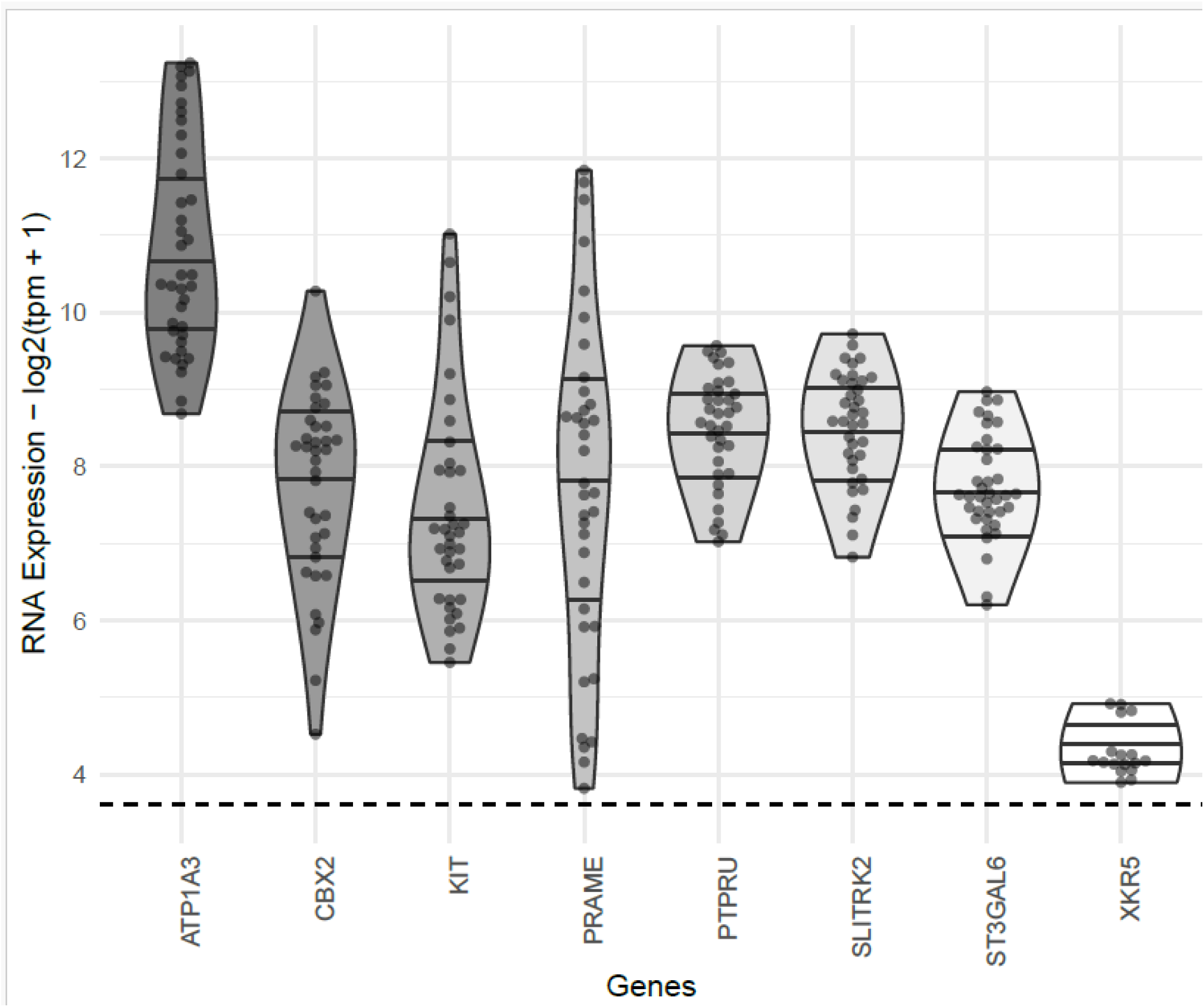
RNA expression of candidate pHLA source genes in DIPG H3K27M autopsy samples. RNA expression values are log2(tpm +1) with variance-stabilising transformation applied from the DESeq2 R package. Expression values for genes in samples for which detected TPM are zero before transformation were not plotted, and a dashed line is shown at this lower limit of detection to aid interpretation.

## Notes

### Competing Interest Statement

The authors have declared no competing interest.

